# De novo transcriptome of *Taverniera cuneifolia* (Roth) Ali

**DOI:** 10.1101/2022.01.27.477879

**Authors:** Talibali Momin, Apurva Punvar, Harshvardhan Zala, Garima Ayachit, Madhvi Joshi, Padamnabhi Nagar

## Abstract

*Taverniera cuneifolia* has been described as a potent substitute of Licorice in India. It has been used as an expectorant, anti-inflammatory, anti-ulcer, wound healing, blood purifier etc. Glycyrrhizin is one of the most useful bioactive sesquiterpenoid present in this plant. The present study aim to carry out transcriptome analysis in root tissue of *Taverniera cuneifolia* to identify specific functional genes involved in the biosynthesis of secondary metabolites. The root transcriptome sequencing of *Taverniera cuneifolia* resulted in a total of ~7.29 Gb of raw data and generated 55,991,233 raw reads. The high quality reads were *de novo* assembled by Trinity assembler followed through CD-HIT resulted into 35,590 “Unigene” transcripts with an average size of 419 bp. The unigenes were analyzed using BLAST2GO resulted in 27,884 (78.35%) transcript with blast hits, 22,510 (63.25%) transcript with mapping and 21,066 (59.19%) transcript with annotation. Functional annotation was carried out using NCBI’s non-redundant and Uniprot databases resulted in the identification of 21,066 (59.19%) annotated transcripts and GO assigned to 24751 (69.54%) transcripts. The gene ontology result shows maximum sequences match with Biological Processes (48%), Molecular Function (27%) and Cellular components (23%). A total of 289 metabolic enriched pathways were identified, which included pathways like Sesquiterpenoid and triterpenoid pathway which were involved in synthesis of secondary metabolite Glycyrrhizin biosynthesis. The enzymes, squalene monooxygenase, farnesyl-diphosphate farnesyltransferase, beta amyrin synthase, beta-amyrin 24-hydroxylase, were identified by functional annotation of transcriptome data. There were several other pathways like terpenoid backbone biosynthesis, steroid biosynthesis, Carotenoid biosynthesis, Flavonoids biosynthesis etc. which have been reported first time from this plant. Transcription factors were predicted by comparison with Plant Transcription Factor Database, and 1557 trancripts belonging to 85 trancription factor families were identified. This transcriptome analysis provided an important resource for future genomic studies in *Taverniera cuneifolia*, therefore representing basis in further investigation of the plant.

**Significance:** Licorice (*Glycyrrhiza glabra* roots) is used as traditional Chinese herbal medicines in majority of formulations. Licorice is also used in Industries like food, herbal and cosmetics etc. due to its high demand in the market it is imported from foreign countries and is not available locally of superior quality (Liu et al., 2015). In India, *Taverniera cuneifolia* has been described as a potent substitute of Licorice, it has been quoted in ancient books like Charak Samhita during the Nigandu period (Kamboj, 2000) and Barda dungar ni Vanaspati ane upyog (Thaker 1910). It has been used as an expectorant, anti-inflammatory, anti-ulcer, wound healing, blood purifier etc. Transcriptomic studies will assist in understanding the basic molecular structure, function and organization of information within the genome of *Taverniera cuniefolia.* This study will help us to identify the key metabolites their expressions and genes responsible for their production.

## 1. Introduction

India is rich in many potential medicinal plants, *Glycyrrhiza glabra* popularly known as Liquorice has been used in the traditional formulation. A licorice (*Glycyrrhiza glabra*) root has been used in more than 1200 formulations in traditional Chinese herbal medicines as major formulations. There are many essential uses of this plant in industries like food, herbal, cosmetics, nutraceuticals etc. (Pastorino et al., 2018). Due to its high demand in the market, it is imported from foreign countries and not available locally of superior quality. In India, *Taverniera cuneifolia* has been described as a potent substitute for Licorice. Glycyrrhizin is one of the most useful bioactive sesquiterpenoid present in this plant.

*Taverniera cuneifolia* belong to fabaceae family, the third largest family of flowering plants, with over 800 genera and 20,000 species. The three major subfamilies include Mimosaceae, Papilionaceae and Caesalpiniacea. The pea (*Pisum sativum* L.) was the model organism used in Mendel’s discovery (1866) and is the foundation of modern plant genetics. The phylogenetic differ greatly in their genome size, base chromosome number, ploidy level and reproductive biology. Two legume species in the Galegoid clade, *Medicago truncatula* and *Lotus japonicus*, from Trifolieae and Loteae tribe respectively, were selected as model system of studying legume genomics and biology. There are many other legumes that have been studies like the soybeans, the most widely grown and economically important legume whose genome has been available since 2010.The common bean (*Phaseolus vulgaris*) the most widely grown grain legume whose genome is available since 2014. Many more legumes have been sequenced since (Smýkal, P. et al., 2020).

*Taverniera cuneifolia* is an important traditional medicinal plant of India as mention in Charak Samita in Nigantu period. It is often referred to as Indian licorice having the same sweet taste as of *Glycyrrhiza glabra* (commercial Licorice) (Zore, 2008). The genus *Taverniera* has sixteen different species (Roskov et al., 2006). It is endemic to North-east Africa and South-west Asian countries (Naik, 1998). Licorice is used as important traditional Chinese medicine with many clinical and industrial applications like Food, Herbal medicine, cosmetics etc. (Liu et al. 2015). *Taverniera cuneifolia* locally known as Jethimad is used by the tribal’s of Barda Hills of Jamnagar in Western India (Saurashtra, Gujarat) as a substitute for Licorice or in other words, the Plant itself is considered to be *Glycyrrhiza glabra* (Nagar, 2005). Many pharmacological benefits of the plants have been reported earlier like expectorant, blood purification, anti-inflammatory, wound healing, anti-ulcer and used in treating spleen tumors (Thaker, Manglorkar and Nagar, 2013).

At the Biochemical level, *Taverniera cuneifolia* has shown the presence of alkaloids, flavonoids, tannins, proteins, reducing sugar and saponins. The presence of oil content in the seeds of *Taverniera cuneifolia* showed polyunsaturated fatty acids, monounsaturated fatty acids and saturated fatty acids (Manglorkar, 2016). *Taverniera cuneifolia* has been assessed very less on phytochemical basis there are only few attempts to characterize this plant at molecular level. *Taverniera cuneifolia* has eight numbers of chromosomes (Perveen and Khatoon, 1989). There is limited information on genetic for this plant on NCBI. Fifteen proteins have been reported from this plants which includes ribosomal protein L32, maturase, photosystem 1 assembly protein Ycf4, cytochrome b6/f complex subunit VIII, D1 protein, photosystem 2 protein M, MaturaseK, ribulose-1,5-bisphosphate carboxylase/oxygenase large subunit, Triosephosphate translocator, Phosphogluconate dehydrogenase, UDP-sulfoquinovose synthase, RNA polymerase beta subunit (Liu et al., 2017).

The current investigation was focused on the most valuable secondary metabolite, Glycyrrhizin and other important secondary metabolites. This experiment provides the in-depth characterizations of this plant. Based on the above facts attempts have been made to identify the genes of various metabolic pathways in *Taverniera cuneifolia* through root transcriptome sequencing. The study will give scientific insight into the molecular network of *Taverniera cuneifolia*.

## Materials and Methods

### Plant material and RNA isolation

*Taverniera cuneifolia* plant was collected from Kutch, Gujarat, India (23.7887 N, 68.79580 E) from its natural habitat near the area of Lakhpat. The tissue of the plant, i.e., roots were cleaned with water than with ethanol and stored in RNA later solution (Qiagen) for longer-term storage. It was then shifted to −20°C in the refrigerator. The total RNA was isolated from the root tissues of the Plant using the RNeasy Plant Mini Kit (Qiagen) following the manufacturer’s instructions. The integrity of the RNA was assessed by formaldehyde agarose gel electrophoresis. Total RNA was quantified by using a Qiaxpert (Qiagen), Qubit 2.0 fluorometer (Life Technologies, Carlsbad, CA, USA) and Qiaxcel capillary electrophoresis (Qiagen). RNA integrity number (RIN) was higher than approx. 7.0 for the sample.

### cDNA library preparation and Sequencing

Ribosomal RNA depletion was carried out using a RiboMinus RNA plant kit for RNA-Seq (Life Technologies, C.A). mRNA fragmentation and cDNA library was constructed using an Ion total RNA-Seq kit v2 (Life Technologies, C.A), further purified using AMpure XP beads (Beckman coulter, Brea, CA, USA). The library was enriched on Ion sphere particles using Dynabeads MyOne Streptavidin C1 using standard protocols for the Ion Proton sequencing. The raw transcriptome data have been deposited in the sequence read archive (SRA) NCBI database with the accession number SRR5626167. This Transcriptome Shotgun Assembly project has been deposited at DDBJ/EMBL/GenBank under the accession GJAF00000000.

### RNA-Seq data processing and *De novo* assembly

Quality control of raw sequence reads was filtered to obtain the high-quality clean reads using bioinformatics tools such as FASTQCv.0.11.5 using a minimum quality threshold Q20 (Andrews, 2010). The clean reads were subjected to de novo assembly using the Trinity v2.4.0 (Grabherr et al., 2011) software to recover full-length transcripts. The redundancy of Trinity generated contigs were clustered for removing duplicate reads with 85% identity using CD-HIT v4.6.1 (Li and Godzik, 2006).

### Functional annotation of transcripts and classification

Functional characterization of assembled sequences was done by performing BlastX of contigs against the non-redundant (nr) database, (https://www.ncbi.nlm.nih.gov/) using an e-value cut-off of 1E-5 followed by further annotation was carried out using Blast2GO (Conesa and Gotz, 2005). Gene Ontology (GO) study was used to classify the functions of the predicted coding sequences. The GO classified the functionally annotated coding sequences into three main domains: Biological process (BP), Molecular function (MF) and Cellular component (CC). Using the Kyoto encyclopedia of genes and genomes (KEGG) (Kanehisa and Goto, 2000) pathway maps were determined. Further, KEGG Automated Annotation Server (KAAS) was used for pathway mapping in addition to Blast2GO (Moriya et al., 2007) for assignment and mapping of the coding DNA sequence (CDS) to the biological pathways. KAAS provides functional annotation of genes by BLAST comparison against the manually curated KEGG genes database.

### Identification of transcription factors families

Transcription factors (TFs) were identified using genome-scale protein and nucleic acid sequences by analyzing InterProScan domain patterns in protein sequences with high coverage and sensitivity using PlantTFcat analysis tool (http://plantgrn.noble.org/PlantTFcat/) tool (Dai et al., 2013).

### SSR prediction

Simple sequence repeats (SSRs) were identified using the MISA tool (Microsatellite; http://pgrc.ipk-gatersleben.de/misa/misa.html). We searched for SSRs ranging from mono to hexanucleotide in size. The minimum repeats number 10 for mononucleotide, 6 for Dinucleotide and 5 for trinucleotide to hexanucleotide was set for SSR search. The maximal number of bases interrupting two SSRs in a compound microsatellite is 100 i.e. the minimum distance between two adjacent SSR markers was set 100 bases.

## Results and Discussion

### Transcriptome Sequencing and *De novo* assembly

The total RNA of two root samples along with RIN value more than 7.0, converted to cDNA library using Ion Total RNA-Seq kit v2 (Life Technologies, C.A), further purified using Ampure XP beads (Beckman coulter, Brea, CA, USA). The library was enriched on Ion sphere particles using Myone C1 Dynabeads. A total of 7.29 gb of raw data was generated using standard protocols for the Ion proton sequencing (Table 1). The good quality roots of *Taverniera cuneifolia* were used for the RNA sequencing, and a total of 55,991,233 reads containing 7,286,727,421 bases were generated. The raw reads were subjected to quality check by FastQC tool and the average base quality was above Q20. De novo transcriptome assembly resulted in 36,896 reads assembled and the final assembly of 35,590 unique high-quality reads was prepared using CD-HIT at 85% sequence similarity, with N50 value of 441 bp. The average GC content of 43% and average contigs length of 419.45 bp was obtained for *Taverniera cuneifolia*. The statistics of transcriptome sequencing and assembly generated by Trinity assembler as given (Table 2).

**Table 1:**
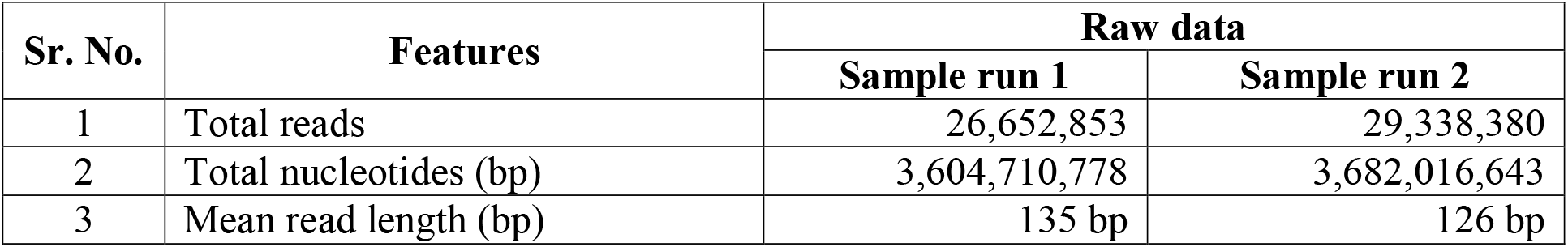
Summary of sequencing data generated for root sample of *Taverniera cuneifolia.*

**Table 2:** Results based on combined assembly of *Taverniera cuneifolia* root transcriptome.

| Sr. No. | Characteristics | Values |
| --- | --- | --- |
| 1 | Total assembled contigs/transcript | 35,590 |
| 2 | GC % | 43.25 |
| 3 | Contig N<sub>50</sub> (bp) | 441 |
| 4 | Median Contig length (bp) | 322 |
| 5 | Average Contig length (bp) | 419.45 |
| 6 | Total assembled bases | 14,928,144 |

### Functional annotation of transcripts

A total of 35,590 transcripts (contigs) assembled by Trinity were subjected to functional annotation using different databases like the Nr Protein database, KEGG, UniProt, etc. GO terms were assigned to transcripts (Supplementary Fig. S2). All transcripts were screened for similarity to a known organism based on the data of species-specific distribution, and it can be concluded that the transcript showed the highest blast hits with *Medicago truncatula* (18,734, 52.63%) followed by *Cicer arietinum* (16,044, 45.08%) and *Glycine max* (15,991, 44.93%). A total of 10590 (29.75%), 8642 (24.28%), 8549 (24.02%), 8399 (23.59%) contigs were found to be similar to *Cajanus cajan*, *Glycine soja*, *Trifolium pratense*, *Trifolium subterraneum*, respectively (Figure 1). The functionally annotated transcripts (27,884, 78.34%) of *Taverniera cuneifolia* were classified using Blast2GO into three main domains; Biological processes, Cellular component and Molecular function gene ontology (Table S1). Among them the most abundant were the Biological processes consisting of 44,395(48.8%) sequences followed by different Molecular Function consisting of 25,025 (27.5%) sequences and last the cellular components consist of 21,508 (23.6%) sequences (Figure 2, 3, 4). The annotated transcripts were subjected to the Kyoto encyclopedia genes and genomes (KEGG) pathway wherein the transcripts were linked to enzymes found in a large number of pathways available in KEGG. The maximum number of annotated transcripts assigned to hydrolases, followed by transferases and oxidoreductases class of enzymes (Figure 5).

**Figure 1:**
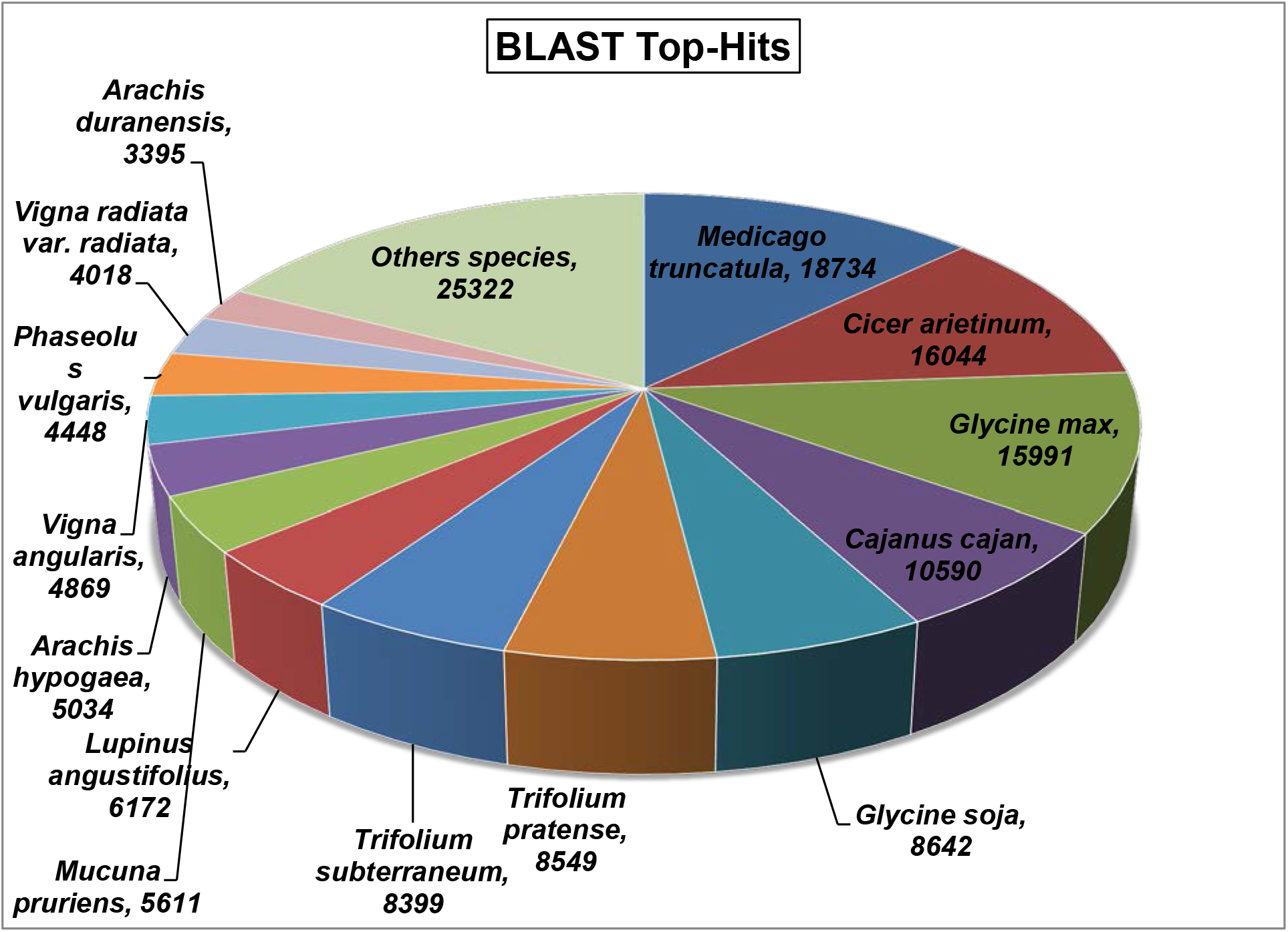
Species distribution of the top BLAST hits of *Taverniera cuneifolia* transcripts in Nr database

**Figure 2:**
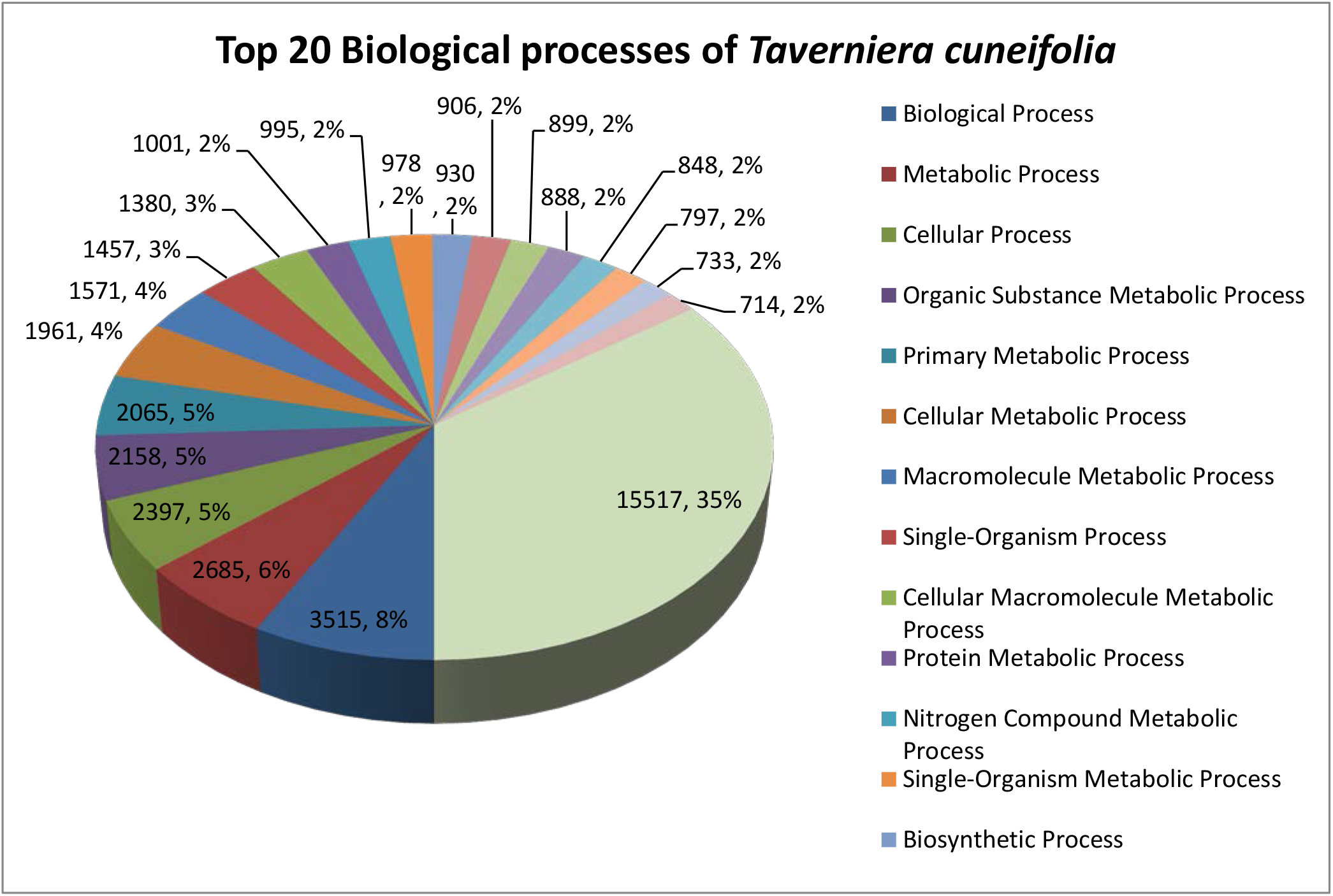
Biological processes gene ontology of *Taverniera cuneifolia* transcripts

**Figure 3:**
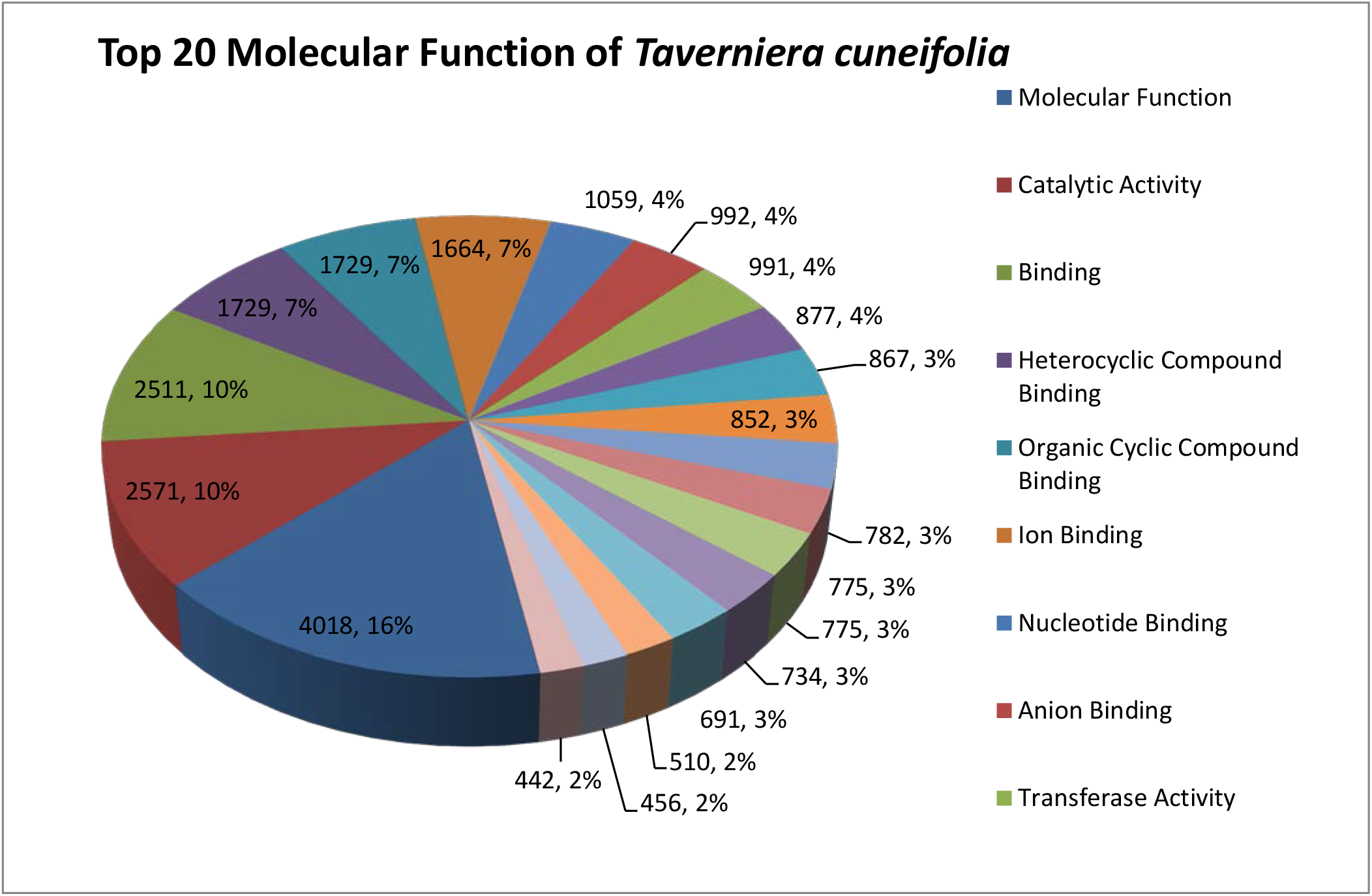
Molecular functions gene ontology of *Taverniera cuneifolia* transcripts

**Figure 4:**
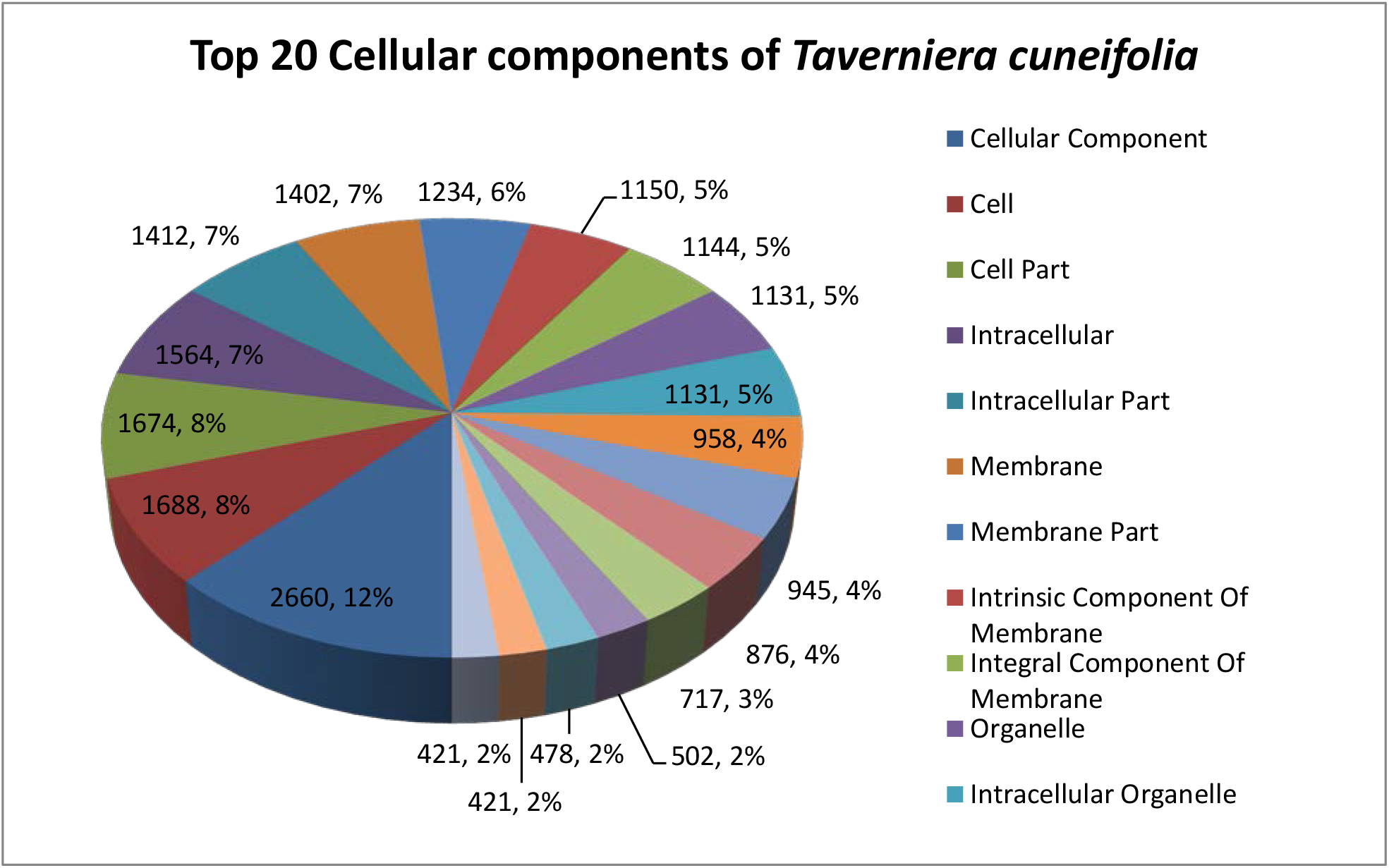
Cellular components gene ontology of *Taverniera cuneifolia* transcripts

**Figure 5:**
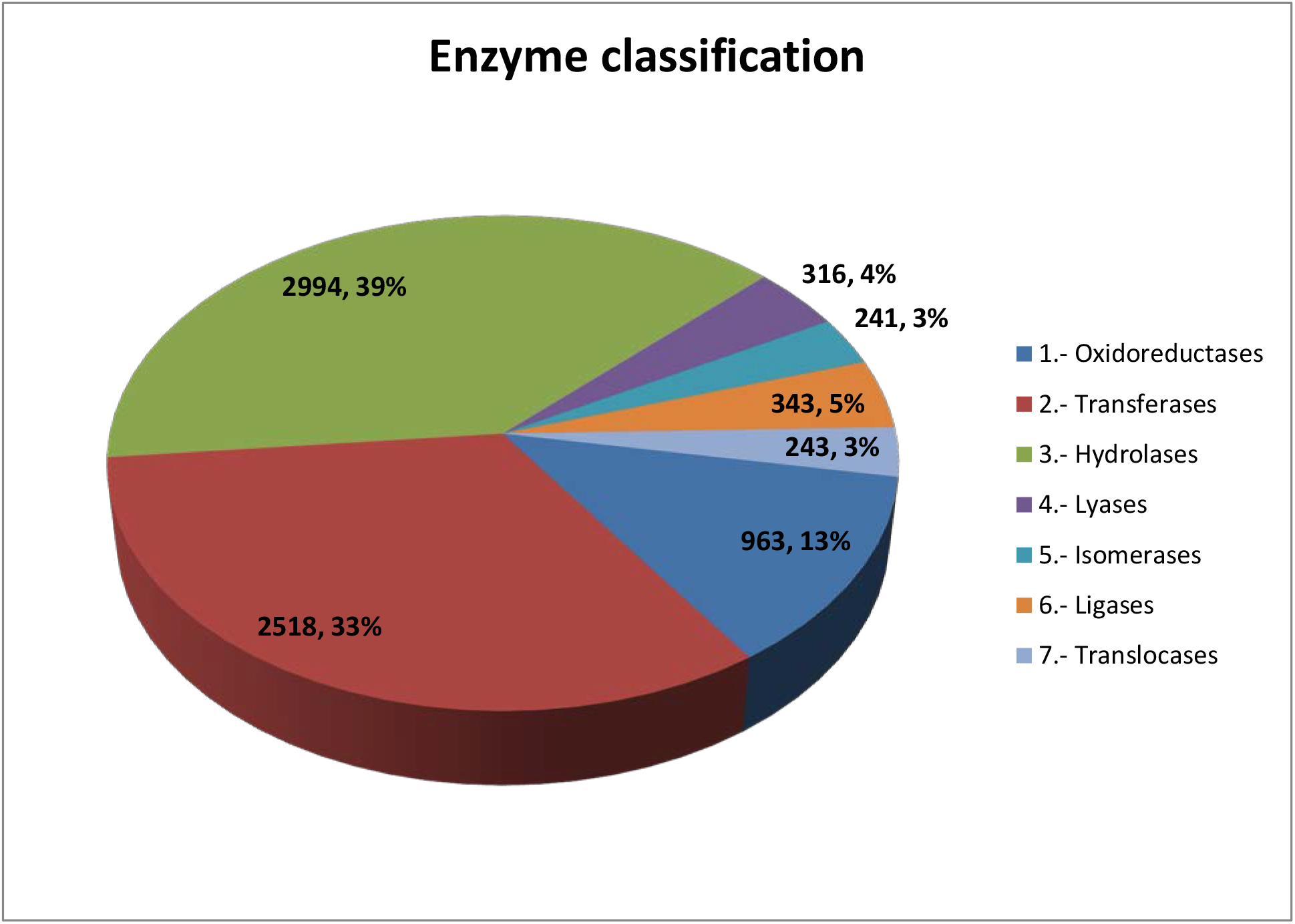
Enzyme classification of *Taverniera cuneifolia* transcripts based on KEGG pathway

### Gene ontology classification

The contigs were further annotated by Blast2Go software with assembled 27,884 transcripts GO terms and divided into three broad categories as Biological Processes (44,395[49%]), Molecular Function (25,025[27%]) and Cellular Component (21,508[24%]) category (Table S1).

The Biological Processes were the most abundant component of GO terms. Among the 44,395 Biological Processes, the maximum number of contigs i.e. represented “Biological process,” followed by “Metabolic process” and “Cellular process” (Figure 2).

A total of 25,025 transcripts were associated with the Molecular function and a relatively large no of the transcript was associated with “Molecular function” followed by “Catalytic activity” and “Binding”, respectively”(Figure 3).

In addition, Cellular Component a total of 21,508 transcripts were associated with the “Cellular component” as the highest match followed by “Cell” and “Cell part” respectively”(Figure 4).

### Pathway Annotation by KEGG

Kyoto Encyclopedia of Genes and Genomes (KEGG) serves as knowledge source to perform functional annotation of the genes. The KEGG represents various biochemical pathways for the genes associated with it. Approximately 289 pathways were annotated and among them, Metabolic pathways (102), Biosynthesis of secondary metabolites (55), Microbial metabolism in diverse environment (22) showed the maximum hit with the database. Some of the important pathways from this plant are discussed below which have been reported with the gene and ko-id. (Table S2).

Terpenoids (isoprenoids) represent the largest and most diverse class of chemicals among the myriad compounds produced by plants. Moreover, the ecological importance of terpenoids has gained increased attention to develop strategies for sustainable pest control and abiotic stress protection. The gene that has shown in this plant includes **Terpenoids backbone biosynthesis (ko00900)** (Supplementary Fig. S3). which includes three gene, ko:K03526 gcpE; (E)-4-hydroxy-3-methylbut-2-enyl-diphosphate synthase [EC: 1.17.7.1 1.17.7.3], ko:K05356 SPS; all-trans-nonaprenyl-diphosphate synthase [EC:2.5.1.84 2.5.1.85], ko:K15889 PCME; prenylcysteine alpha-carboxyl methylesterase [EC:3.1.1.-]. Monoterpenoid biosynthesis having two gene ko: K21373 UGT8; 7-deoxyloganetic acid glucosyltransferase [EC:2.4.1.323], ko:K21374 UGT85A23_24; 7-deoxyloganetin glucosyltransferase [EC:2.4.1.324] and Diterpenoid biosynthesis (ko00904) includes ko:K05282 GA20ox; gibberellin-44 dioxygenase [EC:1.14.11.12].

**Sesquiterpenoid and triterpenoid biosynthesis (ko00909)** (Supplementary Fig. S4). which includes three gene namely ko:K00801 FDFT1; farnesyl-diphosphate farnesyltransferase [EC:2.5.1.21], ko:K15813 LUP4; beta-amyrin synthase [EC:5.4.99.39], ko:K20658 PSM; alpha/beta-amyrin synthase [EC:5.4.99.40 5.4.99.39]. This are the gene on further reactions like oxidation and reductions leads to the production of Glycyrrhizin that is important secondary metabolites as mention above.

**Carotenoid biosynthesis (ko00906)** (Supplementary Fig. S5) includes ko:K09842 AAO3; abscisic-aldehyde oxidase [EC:1.2.3.14], ko:K09843 CYP707A; (+)-abscisic acid 8’-hydroxylase [EC: 1.14.14.137], ko:K14595 AOG; abscisate beta-glucosyltransferase [EC:2.4.1.263].

**Ubiquinone and other terpenoid-quinone biosynthesis (ko00130)** (Supplementary Fig. S6) include ko:K03809 wrbA; NAD(P)H dehydrogenase (quinone) [EC:1.6.5.2].

**Zeatin biosynthesis (ko00908)** (Supplementary Fig. S7) includes ko:K00791 miaA; tRNA dimethylallyltransferase [EC:2.5.1.75], ko:K13496 UGT73C; UDP-glucosyltransferase 73C [EC:2.4.1.-].

**Flavonoid biosynthesis (ko00941)** (Supplementary Fig. S8) inludes ko:K13065 E2.3.1.133; shikimate O-hydroxycinnamoyltransferase [EC:2.3.1.133].

### Candidate genes involved in biosynthesis pathways

Among the 35,591 transcripts that have been annotated using different database, we have identified six gene that play important role in the biosynthesis pathway of Glycyrrhizin production from *Taverniera cuneifolia* (Table S3). Each six different gene includes in formation of Glycyrrhizin.

There were 4912 unigenes hypothetical protein predicted from this plant, of which 30 unigenes that had a hit length above 400 were noted (Table S4). 94 unigenes that predicted Cytochrome P450 family protein from this plant, of which 17 unigenes with a hit length above 150 were noted (Table S5).

## Discussion

Secondary metabolites have key role in providing the defense mechanism to plants against stresses and these metabolites have very important role in many economic important like industries, pharma sector etc (Pagare et al., 2015). There has been no molecular data recorded for this plant as such. The new advancement in the field of omics technologies has led to high-throughtput sequencing data which lead us to prediction of genes, enzymes, complex pathways. (Metzker,2010). De novo of many medicinally important plants such as *Saussurea lappa* (Bains, S et al, 2018), *Vigna radiate* L (Chen, H et al, 2015), *Glycyrrhiza glabra* (Chin,Y et al, 2007), pigeonpea *Cajanus cajan* (L.) Millspaugh (Dutta, S. et al, 2011), *Dracocephalum tanguticum* (Li, H., Fu, Y., Sun, H., Zhang, Y., & Lan, X., 2017) etc. have reported the trancripts involved in active metabolite production using NGS technology.

Transcriptome analysis has proved to be one of the advanced methods for the identification of gene expressing in different pathways of metabolism, growth, development, response towards stress, cell signaling etc. This has help in classifying and categorization different role in secondary metabolic compound. Glycyrrhizin, a well-known secondary metabolite that is found in roots of Licorice has same property that is been found in the roots *Taverniera cuneifolia* which has many uses as described above. A whole transcriptome analysis of root of *Taverniera cuneifolia* has opened the unique transcripts which are reported first time from this plant to be involved in the pathways of primary and secondary metabolism (Sharma, Kumar, Beriwal, et al, 2019).

The de novo assembled transcripts of *Taverniera cuneifolia* were mapped to non-redundant protein database using blastx tool. A total of 35,590 transcripts annotated to the database showed the maximum similarity with *Medicago truncatula* [(18,734) 52.6 %] followed by *Cicer arietinum* [(16,044) 45%] and *Glycine max* [(15,991) 44.9%] and so on, which belong to same family Fabaceae order fabales.

### Main metabolism-related gene of *Taverniera cuneifolia*

Glycyrrhizin is triterpenoid-saponin produced in Licorice roots. It is synthesized via the cytosolic melvonic acid pathway for the production of 2,3-oxidosqualene, which is then cyclized to β-amyrin by β-amyrin synthase (bAS). Then, β-amyrin undergoes a two-step oxidation at the C-30 position followed by glycosylation reactions at the C-3 hydroxyl group to synthesize glycyrrhizin as shown in (Supplementary Fig. S9)(Seki et al 2008, 2011). *Taverniera cuneifolia* also known as Indian Licorice can be used as substitute of *Glycyrrhiza glabra* as it has same features that of this plant. This plant contains varieties of different compound that can be used in future research like triterpenoids, flavonoids, polysaccharides etc, which have been reported first time from this plant. Among them Glycyrrhizin is a primary focus compound that has many economic importance use in different fields. In our experiment we have compared the enzymes and genes for the production of Glycyrrhizin with proposed pathway for biosynthesis of Glycyrrhizin by (Seki et al, 2011), In which Glycyrrhizin is produce by a series of chemical reaction i.e. oxidation of different compound associated with Melvanoic Acid pathway. In this particular pathway there are series of chemical reaction by which Farnesyl diphosphate (FPP) molecule catalyzed by squalene synthase (SQS) originating Squalene. There are fifteen different transcripts that we have found in our plants that are associated for the production of squalene and then by oxidation by squalene epoxidase (SQE) to 2, 3 – oxidosqualene to form β- Amyrin. There are five gene identified from our plant that catalyzed by bAS i.e β- Amyrin synthase to form β- Amyrin. Further β- Amyrin goes into various oxidation reaction with the help of Beta-amyrin 11-oxidase /CYP88D6 and 11-oxo-beta-amyrin 30-oxidase/CYP72A154 to form Glycyrrhetinic acid. The last step includes conversion of glycyrrhetinic acid to glycyrrhizin which includes glycosylation steps in which different enzymes related to UDP-glycuronosyl transferases family are included. There were 32 different UDP-glycuronosyl genes which have been identified from our plant that led to last reaction given in table (Table S3).

At this point *Taverniera cuneifolia* have not been intensively studied and there as such no any reports that showed the details about the enzymes associated in the Glycyrrhizin pathway we have associated with reference pathway proposed by (seki et al 2008, 2011). As there has been no proper investigation for the pathway for glycyrrhizin known till today.

We have extensively worked upon the proteins which we have opted from our data of *Taverniera cuneifolia*. Approx. 4912 genes have been isolated that showed different proteins reported firstly from this plant among them the details have been provided in (Table S4) (we have approx. shown only those hypothetical proteins whose hit length is above 400 bases). In our studies we also found that there were more than 90 transcripts that showed the function related to Cytochrome P450 family protein. This protein has an immense ability to synthesis many new molecules required in the system to function and cope up with.

### Identification of SSR markers and Transcription factors

The potential SSR from mono to hexanucleotide were predicted using MISA Perl script. A total of 35,590 unigene sequences were examined and 2912 SSR were obtained. It was found that only 2454 number of sequences were containing SSRs. Further, only 365 sequences contained >1 SSR marker and 265 were present in compound form. Tri-nucleotide represented the maximum numbers of SSRs (1291), followed by Mono-nucleotide (832) and then Di-nucleotide (597) (Table 3). The analysis of the transcripts revealed 1557 unique transcripts belonging to 85 transcription factor families. Among the identified unigenes, the highest of them represented the WD40 family followed by C2H2, MYB-HB, AP2-EREBP, PHD etc. the top 15 have been shown in the table.(Figure 6).

**Figure 6:**
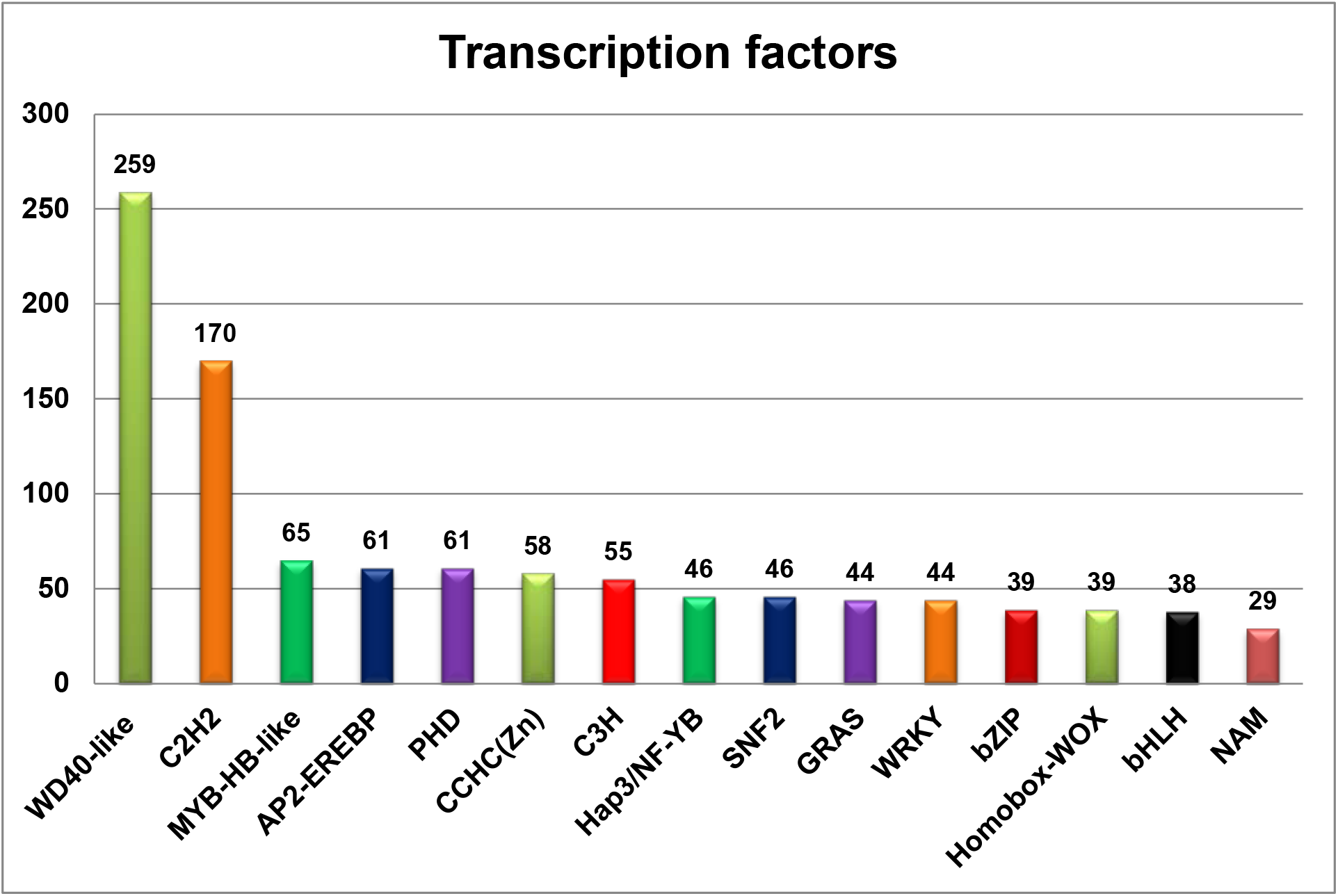
Top 15 Transcription factors families detection from *Taverniera cuneifolia* root transcriptome.

**Table 3:** Identification of Simple Sequence Repeats (SSRs) from *Taverniera cuneifolia* root transcriptome.

| SSR statistics | Count |
| --- | --- |
| Total number of sequences examined | 35,590 |
| Total size of examined sequences (bp) | 1,49,28,144 |
| Total number of identified SSRs | 2,912 |
| Number of SSR containing sequences | 2,454 |
| Number of sequences containing more than 1 SSR | 365 |
| Number of SSRs present in compound formation | 265 |
| Mono-nucleotide | 832 |
| Di-nucleotide | 597 |
| Tri-nucleotide | 1291 |
| Tetra-nucleotide | 153 |
| Penta-nucleotide | 33 |
| Hexa-nucleotide | 6 |

**Table 4:**
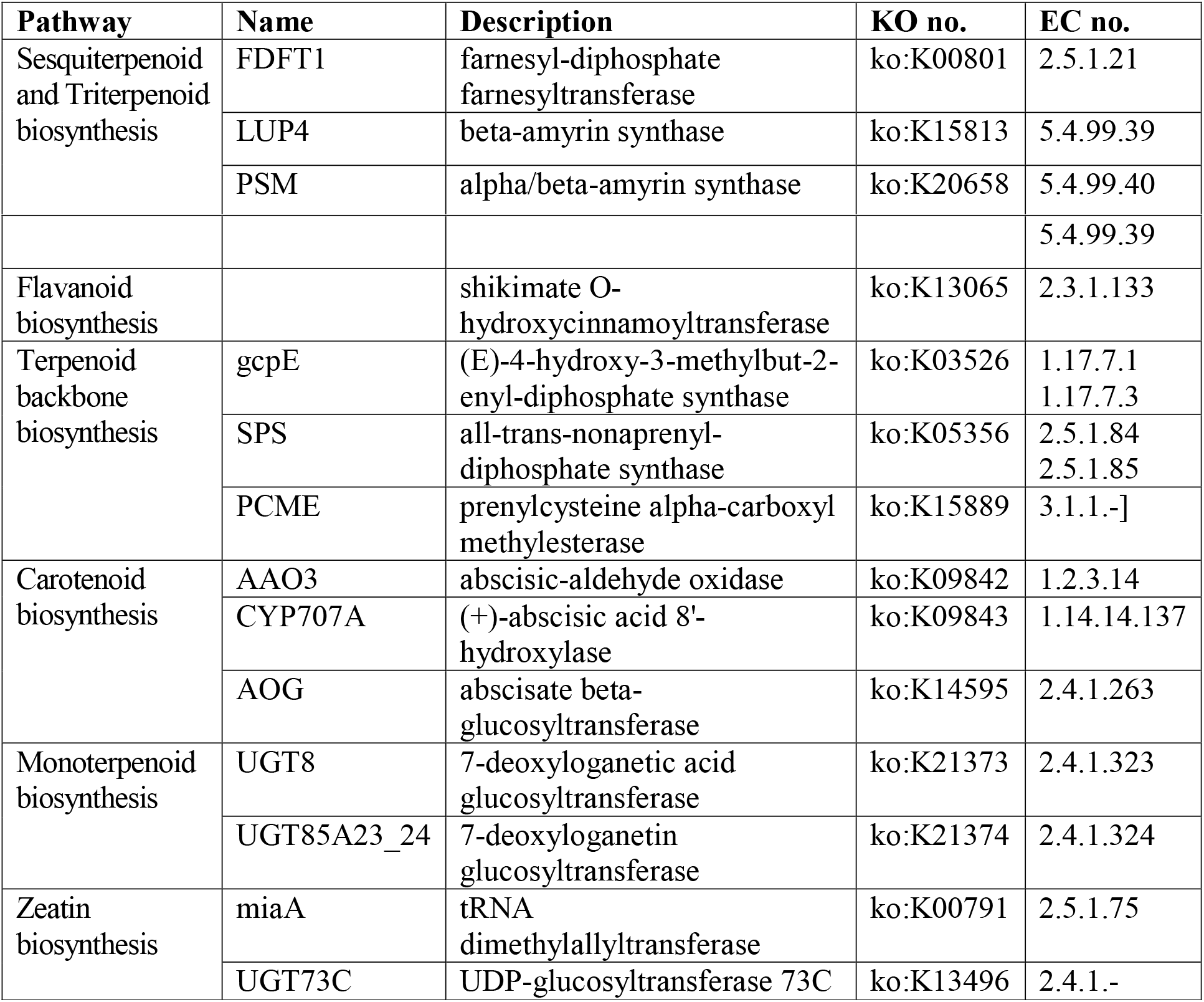
Candidate “Unigenes” encoding enzymes involved in the Sesquiterpenoid and Triterpenoid biosynthesis, Flavonoid biosynthesis, Terpenoid backbone biosynthesis, Carotenoid biosynthesis, Monoterpenoid biosynthesis and Zeatin biosynthesis identified from *Taverniera cuneifolia* Transcriptome.

## Acknowledgments

We are grateful to GBRC (Gujarat Biotechnology Research Centre) for providing the platform for performing the experiment. All the facilities were provided by GBRC including Computational Analysis. The Department of Botany, Faculty of Science, The Maharaja Sayajirao University Baroda for the all the supports for this work.

## Author’s contribution

All authors have contributed to various aspects of this work. PN and MJ conceived the idea and designed the experiments. TM and HZ performed the experiment. TM, HZ, GA and AP analyzed the data. TM analyzed the results and wrote the manuscript. PSN, HZ and MJ finalized the manuscript.

## Supplementary Tables

**Table S1:** GO sequence distribution of biological processes, molecular functions and cellular components.

| GO term | Process | No. of Transcripts |
| --- | --- | --- |
| **Biological processes (44,395)** | Biological process | 3515 |
|  | Metabolic process | 2685 |
|  | Cellular process | 2397 |
|  | Organic substance metabolic process | 2158 |
|  | Primary metabolic process | 2065 |
|  | Cellular metabolic process | 1961 |
|  | Macromolecule metabolic process | 1571 |
|  | Single-organism process | 1457 |
|  | Cellular macromolecule metabolic process | 1380 |
|  | Protein metabolic process | 1001 |
|  | Nitrogen compound metabolic process | 995 |
|  | Single-organism metabolic process | 978 |
|  | Biosynthetic process | 930 |
|  | Organic substance biosynthetic process | 906 |
|  | Cellular nitrogen compound metabolic process | 899 |
|  | Cellular biosynthetic process | 888 |
|  | Cellular protein metabolic process | 848 |
|  | Single-organism cellular process | 797 |
|  | Organic cyclic compound metabolic process | 733 |
|  | Heterocycle metabolic process | 714 |
|  | Cellular aromatic compound metabolic process | 712 |
|  | Macromolecule biosynthetic process | 673 |
|  | Nucleobase-containing compound metabolic process | 659 |
|  | Cellular macromolecule biosynthetic process | 650 |
|  | Cellular nitrogen compound biosynthetic process | 637 |
|  | Phosphorus metabolic process | 632 |
|  | Gene expression | 631 |
|  | Phosphate-containing compound metabolic process | 629 |
|  | Biological regulation | 614 |
|  | Macromolecule modification | 589 |
|  | Protein modification process | 576 |
|  | Cellular protein modification process | 576 |
|  | Regulation of biological process | 555 |
|  | Localization | 533 |
|  | Nucleic acid metabolic process | 531 |
|  | Establishment of localization | 530 |
|  | Transport | 529 |
|  | Regulation of cellular process | 513 |
|  | Organonitrogen compound metabolic process | 493 |
|  | Oxidation-reduction process | 483 |
|  | Phosphorylation | 478 |
|  | RNA metabolic process | 459 |
|  | Organic cyclic compound biosynthetic process | 437 |
|  | Response to stimulus | 433 |
|  | Heterocycle biosynthetic process | 421 |
|  | Aromatic compound biosynthetic process | 416 |
|  | Small molecule metabolic process | 386 |
|  | Nucleobase-containing compound biosynthetic process | 383 |
|  | Organonitrogen compound biosynthetic process | 359 |
| **Cellular components**<br>**(21,508)** | Cellular component | 2660 |
|  | Cell | 1688 |
|  | Cell part | 1674 |
|  | Intracellular | 1564 |
|  | Intracellular part | 1412 |
|  | Membrane | 1402 |
|  | Membrane part | 1234 |
|  | Intrinsic component of membrane | 1150 |
|  | Integral component of membrane | 1144 |
|  | Organelle | 1131 |
|  | Intracellular organelle | 1131 |
|  | Membrane-bounded organelle | 958 |
|  | Intracellular membrane-bounded organelle | 945 |
|  | Cytoplasm | 876 |
|  | Cytoplasmic part | 717 |
|  | Macromolecular complex | 502 |
|  | Nucleus | 478 |
|  | Organelle part | 421 |
|  | Intracellular organelle part | 421 |
| **Molecular functions**<br>**(25,025)** | Molecular function | 4018 |
|  | Catalytic activity | 2571 |
|  | Binding | 2511 |
|  | Heterocyclic compound binding | 1729 |
|  | Organic cyclic compound binding | 1729 |
|  | Ion binding | 1664 |
|  | Nucleotide binding | 1059 |
|  | Transferase activity | 991 |
|  | Ribonucleoside binding | 877 |
|  | Purine ribonucleoside binding | 867 |
|  | Hydrolase activity | 852 |
|  | Adenyl ribonucleotide binding | 775 |
|  | Oxidoreductase activity | 456 |
|  | Anion binding | 992 |
|  | Cation binding | 782 |
|  | ATP binding | **734** |
|  | Nucleic acid binding | **691** |
|  | Transferase activity, transferring phosphorus-containing groups | **510** |
|  | Metal ion binding | **775** |
|  | Kinase activity | **442** |

**Table S2:** Distribution of transcripts to biological pathways using KEGG specific to plants along with KO-ID.

| **KO-ID** | **KEGGS Pathways Distribution** | **Transcripts no.** |
| --- | --- | --- |
| ko01100 | Metabolic pathways | 102 |
| ko01110 | Biosynthesis of secondary metabolites | 55 |
| ko01120 | Microbial metabolism in diverse environments | 22 |
| ko04075 | Plant hormone signal transduction | 15 |
| ko03040 | Spliceosome | 14 |
| ko01200 | Carbon metabolism | 13 |
| ko03010 | Ribosome | 13 |
| ko04144 | Endocytosis | 13 |
| ko03013 | RNA transport | 12 |
| ko04714 | Thermogenesis | 12 |
| ko04626 | Plant-pathogen interaction | 12 |
| ko04016 | MAPK signaling pathway - plant | 11 |
| ko04120 | Ubiquitin mediated proteolysis | 10 |
| ko04141 | Protein processing in endoplasmic reticulum | 10 |
| ko01230 | Biosynthesis of amino acids | 10 |
| ko00520 | Amino sugar and nucleotide sugar metabolism | 10 |
| ko03018 | RNA degradation | 10 |
| ko00190 | Oxidative phosphorylation | 9 |
| ko03015 | mRNA surveillance pathway | 8 |
| ko00500 | Starch and sucrose metabolism | 7 |
| ko01240 | Biosynthesis of cofactors | 7 |
| ko04146 | Peroxisome | 7 |
| ko00620 | Pyruvate metabolism | 7 |
| ko00010 | Glycolysis / Gluconeogenesis | 6 |
| ko03050 | Proteasome | 5 |
| ko00052 | Galactose metabolism | 5 |
| ko04810 | Regulation of actin cytoskeleton | 5 |
| ko00051 | Fructose and mannose metabolism | 5 |
| ko03008 | Ribosome biogenesis in eukaryotes | 5 |
| ko00270 | Cysteine and methionine metabolism | 5 |
| ko00240 | Pyrimidine metabolism | 5 |
| ko04142 | Lysosome | 5 |
| ko00920 | Sulfur metabolism | 4 |
| ko00410 | beta-Alanine metabolism | 4 |
| ko03420 | Nucleotide excision repair | 4 |
| ko00230 | Purine metabolism | 4 |
| ko00983 | Drug metabolism - other enzymes | 4 |
| ko04110 | Cell cycle | 4 |
| ko00020 | Citrate cycle | 4 |
| ko00970 | Aminoacyl-tRNA biosynthesis | 4 |
| ko04072 | Phospholipase D signaling pathway | 4 |
| ko00250 | Alanine, aspartate and glutamate metabolism | 4 |
| ko00720 | Carbon fixation pathways in prokaryotes | 4 |
| ko00940 | Phenylpropanoid biosynthesis | 4 |
| ko00564 | Glycerophospholipid metabolism | 4 |
| ko00710 | Carbon fixation in photosynthetic organisms | 4 |
| ko00480 | Glutathione metabolism | 4 |
| ko01212 | Fatty acid metabolism | 4 |
| ko00906 | Carotenoid biosynthesis | 3 |
| ko00030 | Pentose phosphate pathway | 3 |
| ko00071 | Fatty acid degradation | 3 |
| ko00280 | Valine, leucine and isoleucine degradation | 3 |
| ko00330 | Arginine and proline metabolism | 3 |
| ko04712 | Circadian rhythm - plant | 3 |
| ko00040 | Pentose and glucuronate interconversions | 3 |
| ko01210 | 2-Oxocarboxylic acid metabolism | 3 |
| ko03020 | RNA polymerase | 3 |
| ko00909 | Sesquiterpenoid and triterpenoid biosynthesis | 3 |
| ko00900 | Terpenoid backbone biosynthesis | 3 |
| ko00460 | Cyanoamino acid metabolism | 2 |
| ko03410 | Base excision repair | 2 |
| ko00902 | Monoterpenoid biosynthesis | 2 |
| ko00340 | Histidine metabolism | 2 |
| ko00908 | Zeatin biosynthesis | 2 |
| ko00980 | Metabolism of xenobiotics by cytochrome P450 | 2 |
| ko03440 | Homologous recombination | 2 |
| ko00061 | Fatty acid biosynthesis | 2 |
| ko00630 | Glyoxylate and dicarboxylate metabolism | 2 |
| ko02010 | ABC transporters | 2 |
| ko01040 | Biosynthesis of unsaturated fatty acids | 2 |
| ko04978 | Mineral absorption | 2 |
| ko04024 | cAMP signaling pathway | 2 |
| ko00592 | alpha-Linolenic acid metabolism | 2 |
| ko00290 | Valine, leucine and isoleucine biosynthesis | 2 |
| ko00260 | Glycine, serine and threonine metabolism | 2 |
| ko00625 | Chloroalkane and chloroalkene degradation | 2 |
| ko00510 | N-Glycan biosynthesis | 2 |
| ko04310 | Wnt signaling pathway | 2 |
| ko04020 | Calcium signaling pathway | 2 |
| ko00310 | Lysine degradation | 2 |
| ko03060 | Protein export | 2 |
| ko04922 | Glucagon signaling pathway | 2 |
| ko00640 | Propanoate metabolism | 2 |
| ko00941 | Flavonoid biosynthesis | 1 |
| ko03430 | Mismatch repair | 1 |
| ko00220 | Arginine biosynthesis | 1 |
| ko02020 | Two-component system | 1 |
| ko00945 | Stilbenoid, diarylheptanoid and gingerol biosynthesis | 1 |
| ko00513 | Various types of N-glycan biosynthesis | 1 |
| ko00903 | Limonene and pinene degradation | 1 |
| ko00966 | Glucosinolate biosynthesis | 1 |
| ko00730 | Thiamine metabolism | 1 |
| ko00400 | Phenylalanine, tyrosine and tryptophan biosynthesis | 1 |
| ko04122 | Sulfur relay system | 1 |
| ko00100 | Steroid biosynthesis | 1 |
| ko00380 | Tryptophan metabolism | 1 |
| ko00982 | Drug metabolism - cytochrome P450 | 1 |
| ko00790 | Folate biosynthesis | 1 |
| ko03030 | DNA replication | 1 |
| ko04979 | Cholesterol metabolism | 1 |
| ko00904 | Diterpenoid biosynthesis | 1 |
| ko03022 | Basal transcription factors | 1 |
| ko00130 | Ubiquinone and other terpenoid-quinone biosynthesis | 1 |

**Table S3:**
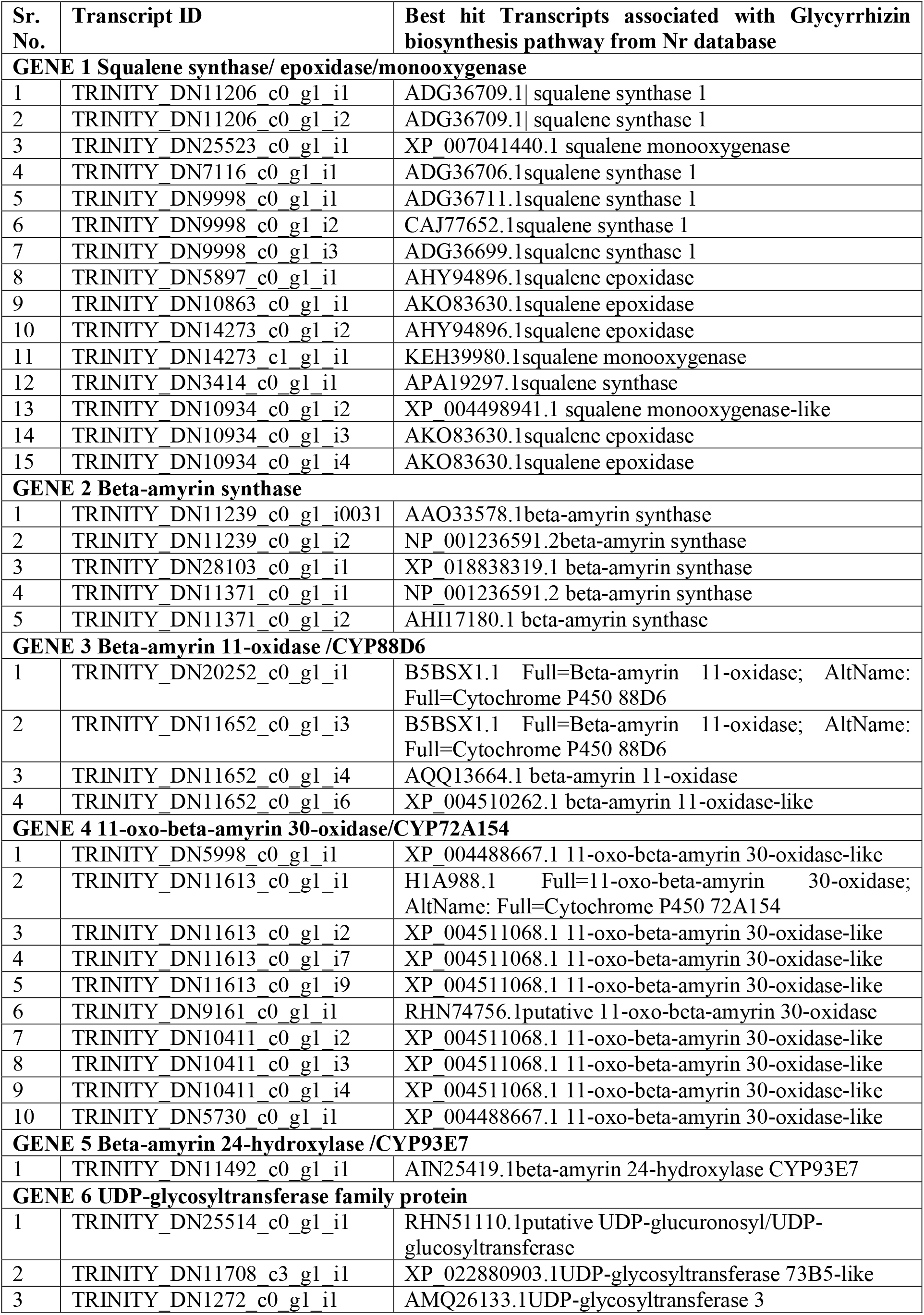
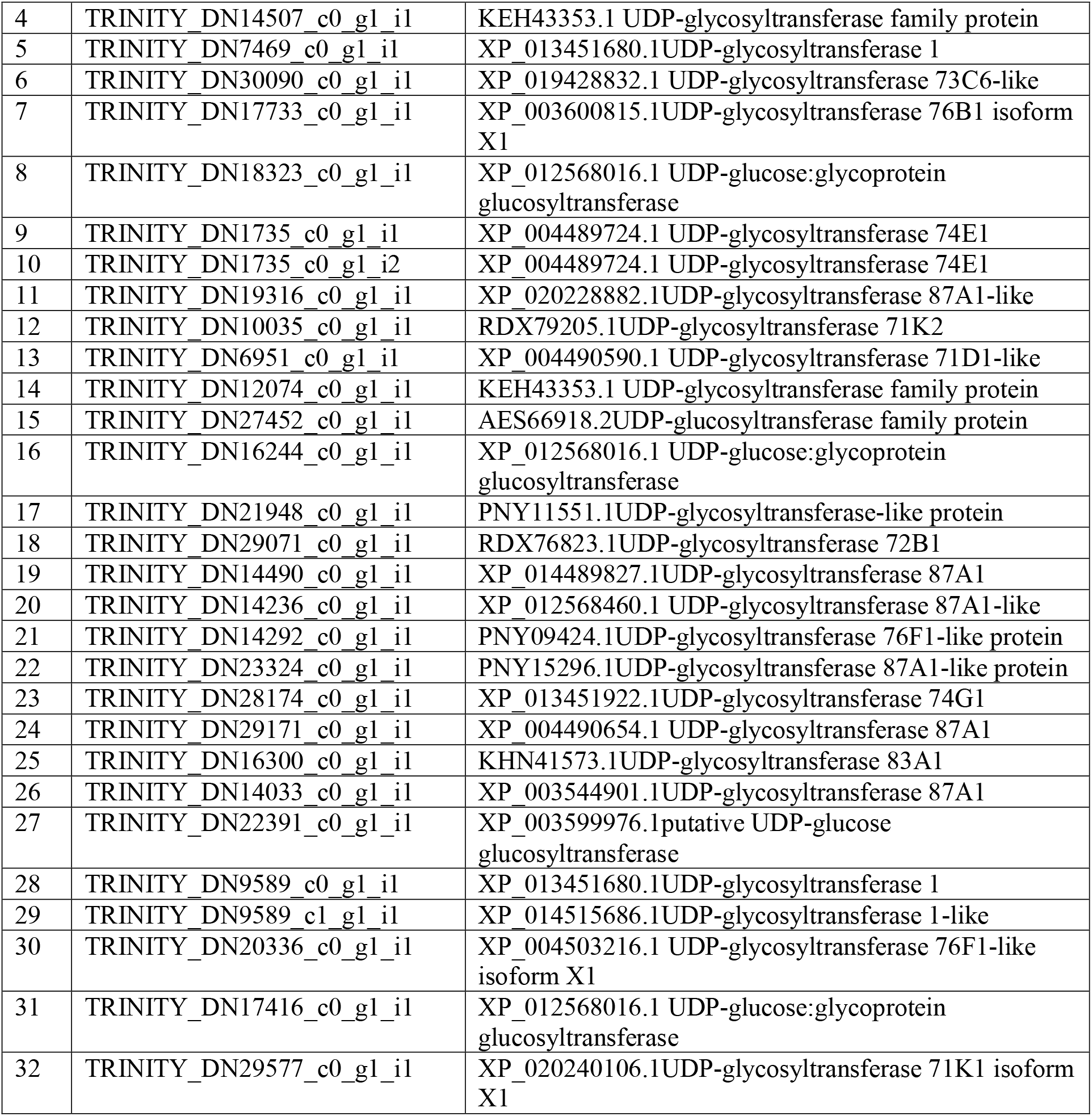
Transcripts/genes that is associated with *Glycyrrhizin* production in *Taverniera cuneifolia* from Nr database.

**Table S4:**
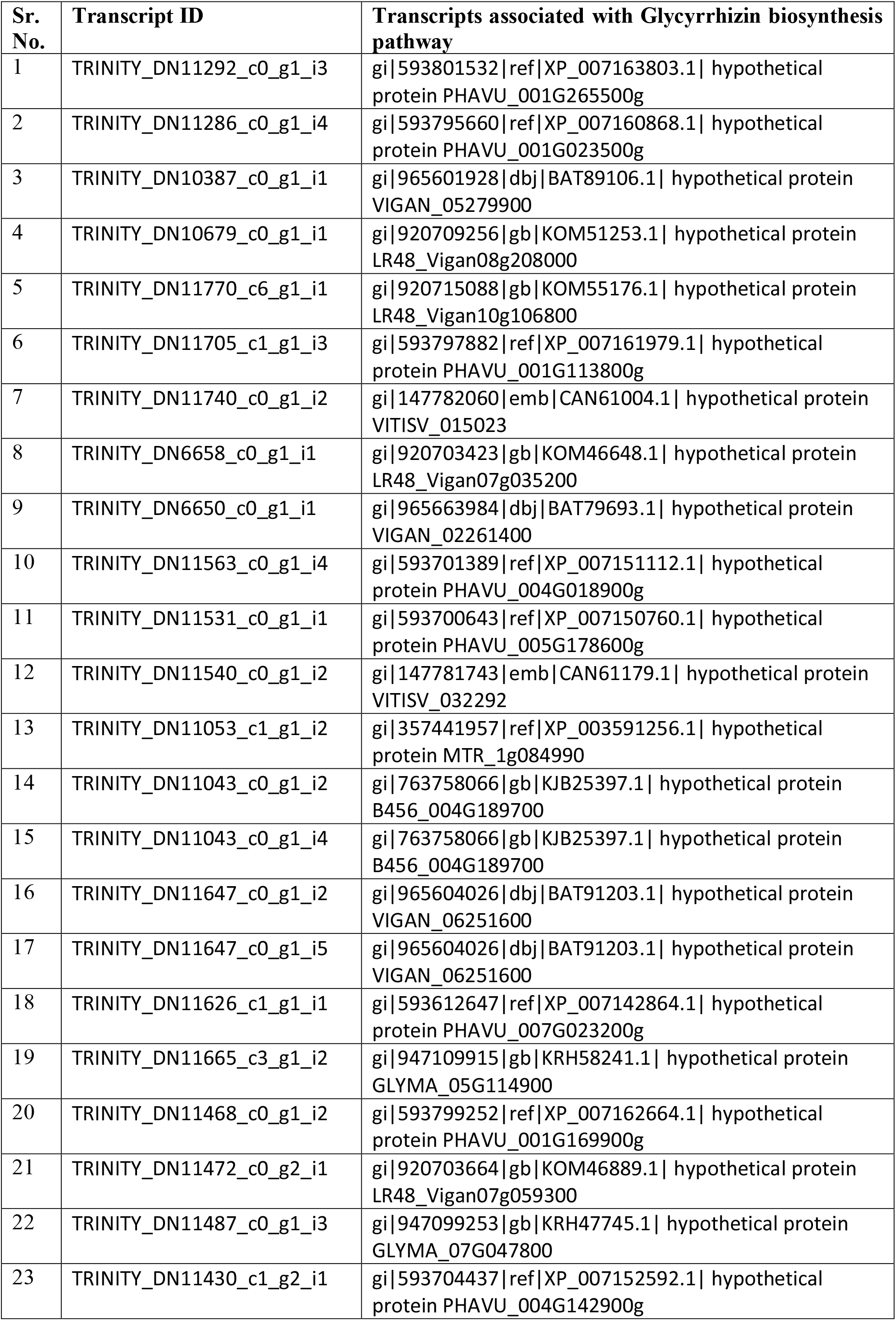
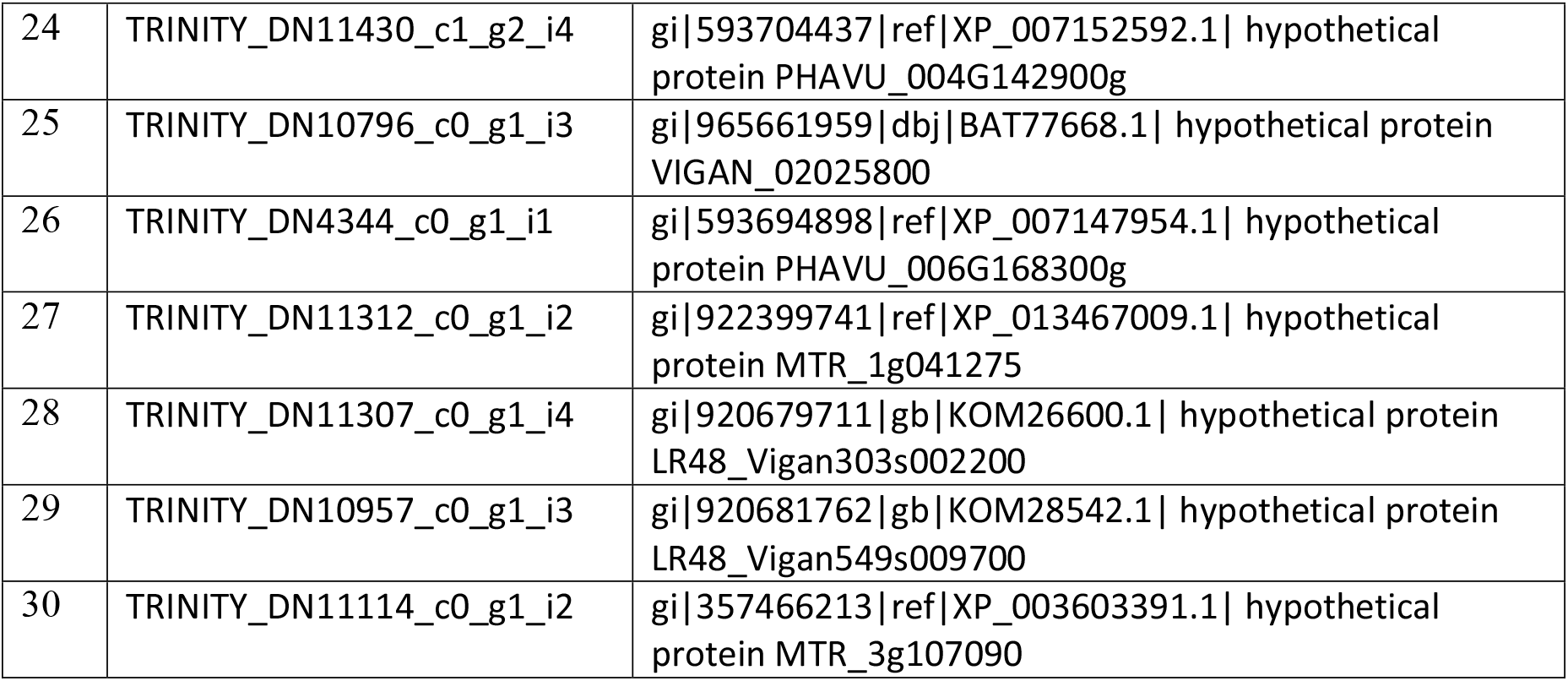
Transcripts/genes that showed the Hypothetical protein in *Taverniera cuneifolia* with hit length above 400, (total over all 4912 hypothetical protien).

**Table S5:** Transcripts/genes that showed the Cytochrome P450 family protein in *Taverniera cuneifolia*

| Sr. No. | Transcript ID | Transcripts associated with Cytochrome P450 family protein |
| --- | --- | --- |
| 1 | TRINITY_DN10399_c0_g1_i2 | gi\|356515730\|ref\|XP_003526551.1\| PREDICTED: NADPH--cytochrome P450 reductase |
| 2 | TRINITY_DN11569_c1_g1_i1 | gi\|357514033\|ref\|XP_003627305.1\| cytochrome P450 family monooxygenase |
| 3 | TRINITY_DN11569_c1_g1_i3 | gi\|357514033\|ref\|XP_003627305.1\| cytochrome P450 family monooxygenase |
| 4 | TRINITY_DN1922_c0_g1_i2 | gi\|502156756\|ref\|XP_004510631.1\| PREDICTED: cytochrome P450 78A3 |
| 5 | TRINITY_DN11093_c0_g1_i2 | gi\|922394449\|ref\|XP_013465628.1\| cytochrome P450 family Ent-kaurenoic acid oxidase |
| 6 | TRINITY_DN11010_c1_g1_i1 | gi\|357470373\|ref\|XP_003605471.1\| cytochrome P450 family monooxygenase |
| 7 | TRINITY_DN11652_c0_g1_i4 | gi\|838228579\|gb\|AKM97308.1\| cytochrome P450 88D6 |
| 8 | TRINITY_DN8146_c0_g1_i1 | gi\|371940464\|dbj\|BAL45206.1\| cytochrome P450 monooxygenase |
| 9 | TRINITY_DN9931_c0_g1_i2 | gi\|502150242\|ref\|XP_004507858.1\| PREDICTED: NADPH--cytochrome P450 reductase |
| 10 | TRINITY_DN9161_c0_g1_i1 | gi\|922392052\|ref\|XP_013464437.1\| cytochrome P450 family protein |
| 11 | TRINITY_DN3071_c0_g1_i1 | gi\|922399435\|ref\|XP_013466867.1\| cytochrome P450 family protein |
| 12 | TRINITY_DN5499_c0_g1_i1 | gi\|922394449\|ref\|XP_013465628.1\| cytochrome P450 family Ent-kaurenoic acid oxidase |
| 13 | TRINITY_DN2780_c0_g1_i1 | gi\|502161259\|ref\|XP_004512097.1\| PREDICTED: cytochrome P450 84A1 |
| 14 | TRINITY_DN2767_c0_g1_i1 | gi\|356569428\|ref\|XP_003552903.1\| PREDICTED: cytochrome P450 714C2-like |
| 15 | TRINITY_DN10453_c0_g1_i1 | gi\|356540462\|ref\|XP_003538708.1\| PREDICTED: cytochrome P450 87A3-like |
| 16 | TRINITY_DN10993_c0_g1_i1 | gi\|922380457\|ref\|XP_013460458.1\| cytochrome P450 family protein |
| 17 | TRINITY_DN11158_c1_g1_i4 | gi\|922327835\|ref\|XP_013443310.1\| cytochrome P450 family 71 protein |

## Supplementary Figures

**Supplementary Fig. S1:**
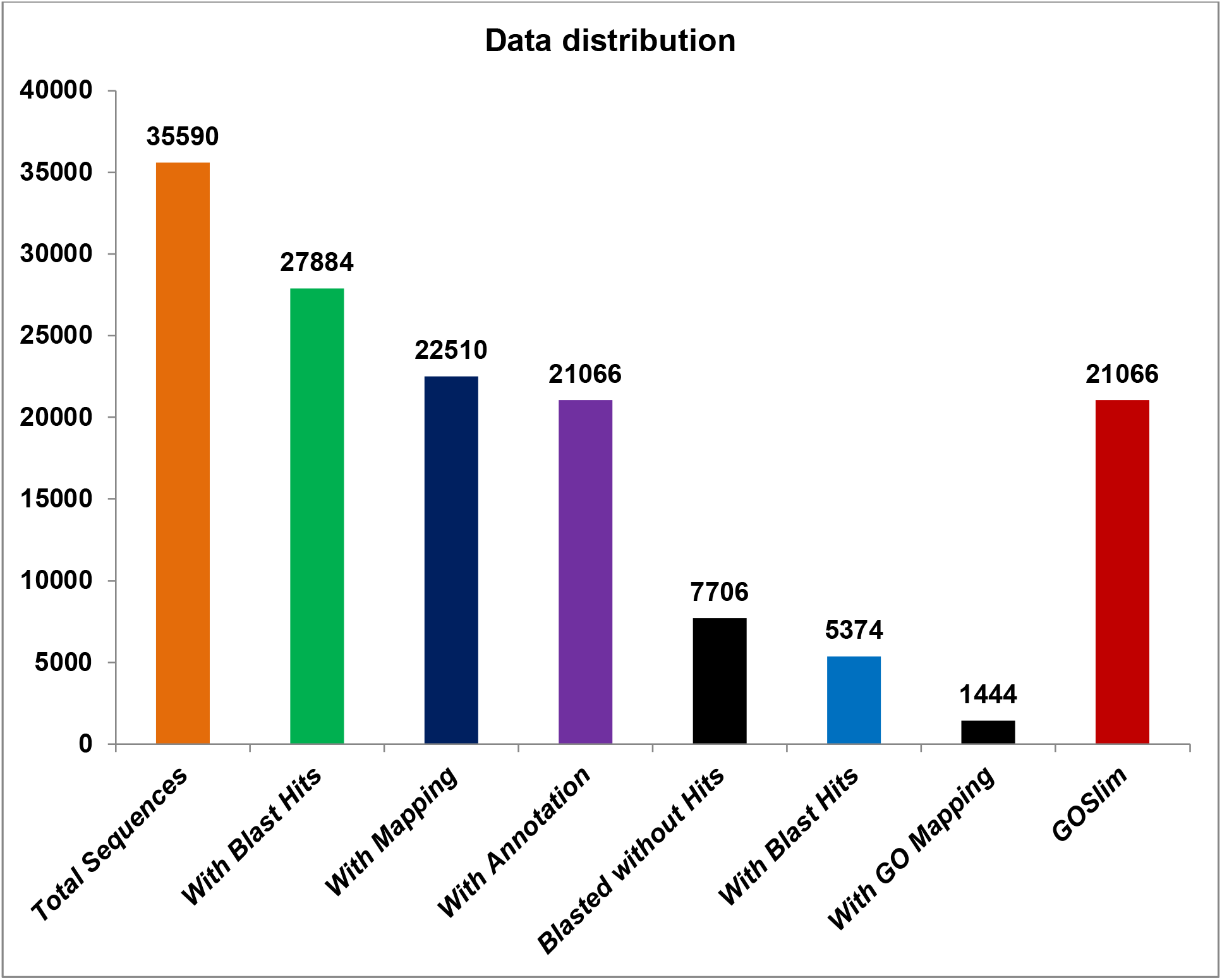
Data distribution of *Taverniera cuneifolia* transcripts subjected to functional annotation.

**Supplementary Fig. S2:**
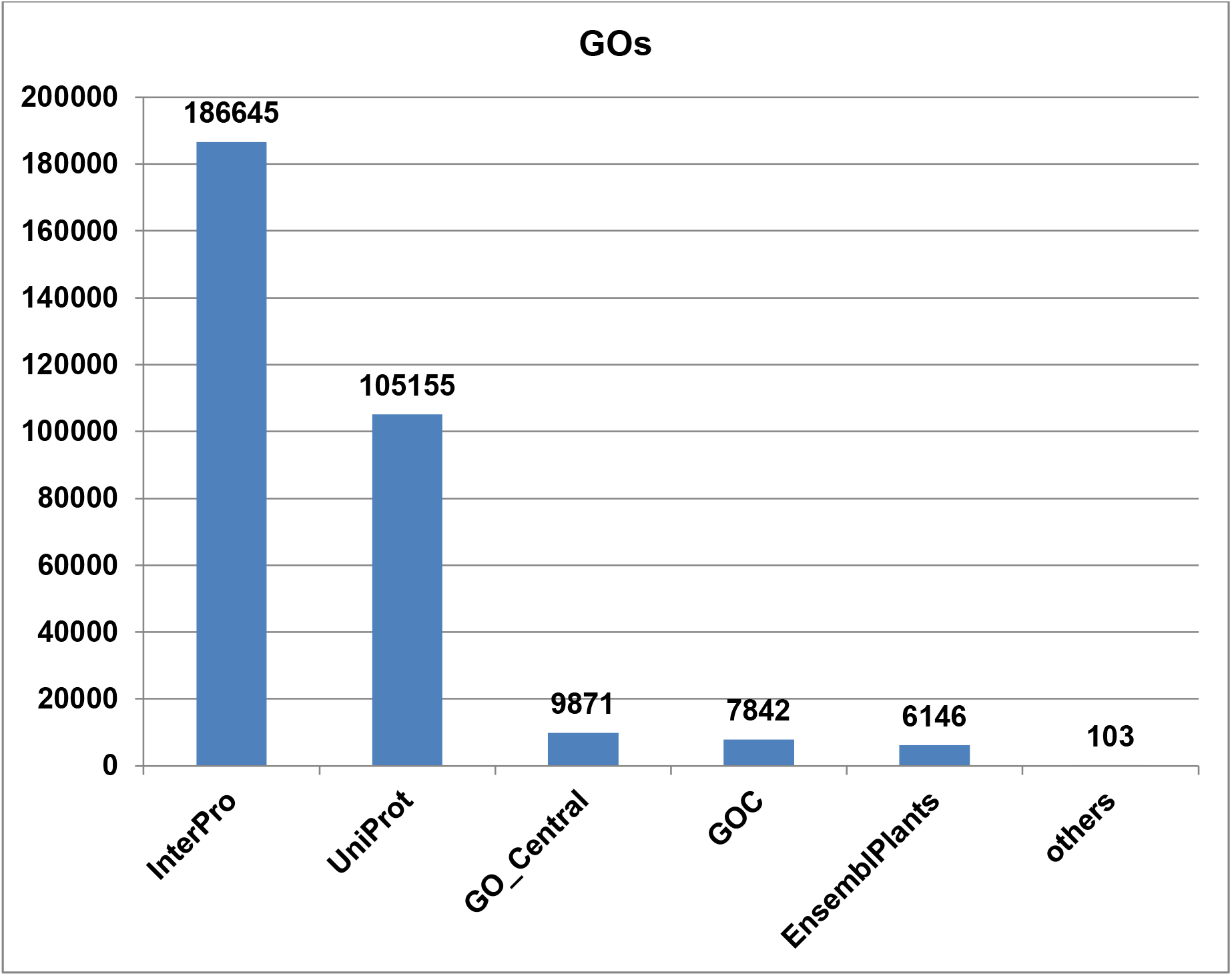
Annotation of *Taverniera cuneifolia* transcripts to different database sources.

**Supplementary Fig. S3:**
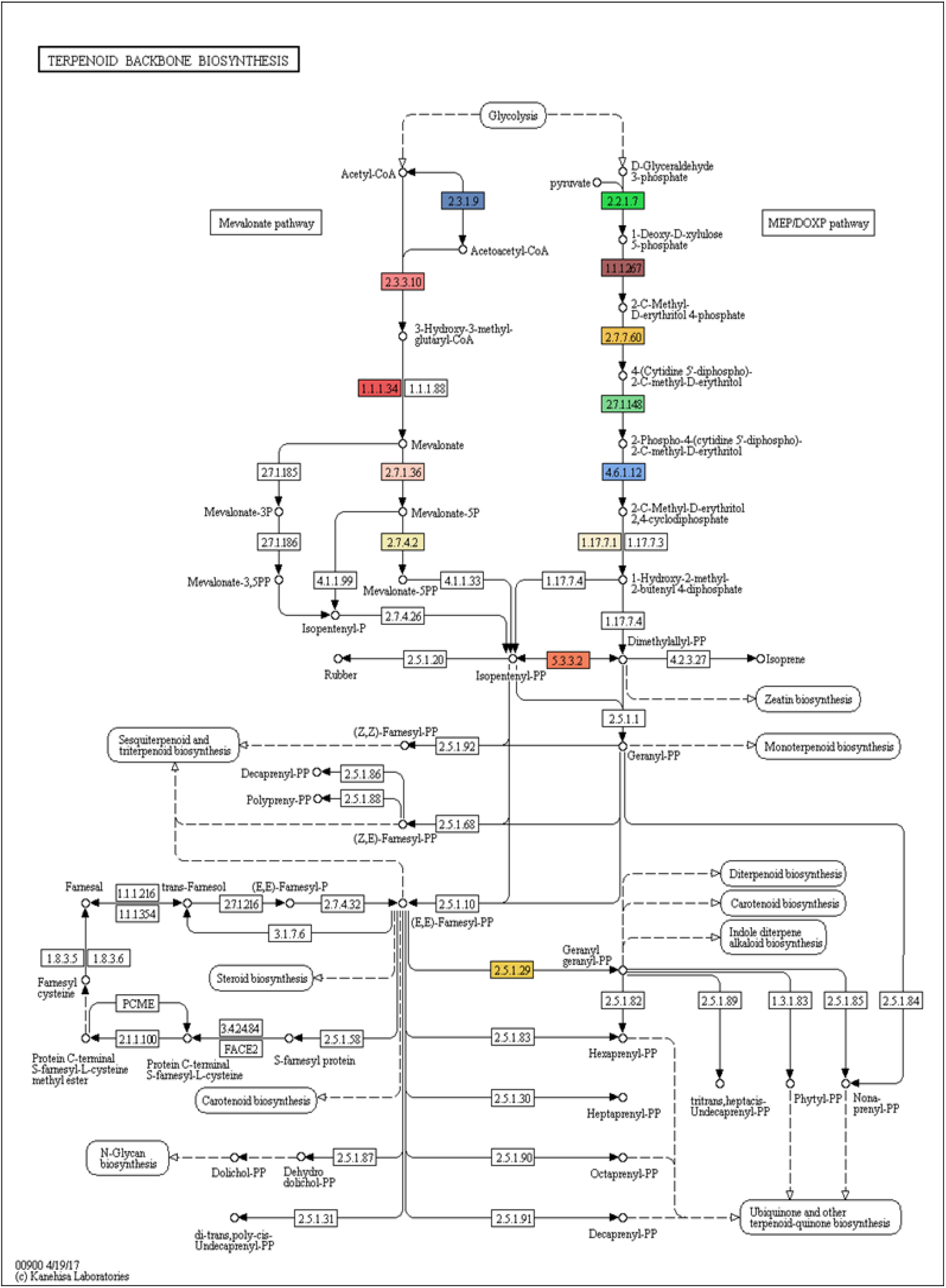
Terpenoids backbone biosynthesis pathways (Ko00900), color boxes are the gene found from *Taverniera cuneifolia* sequences.

**Supplementary Fig. S4:**
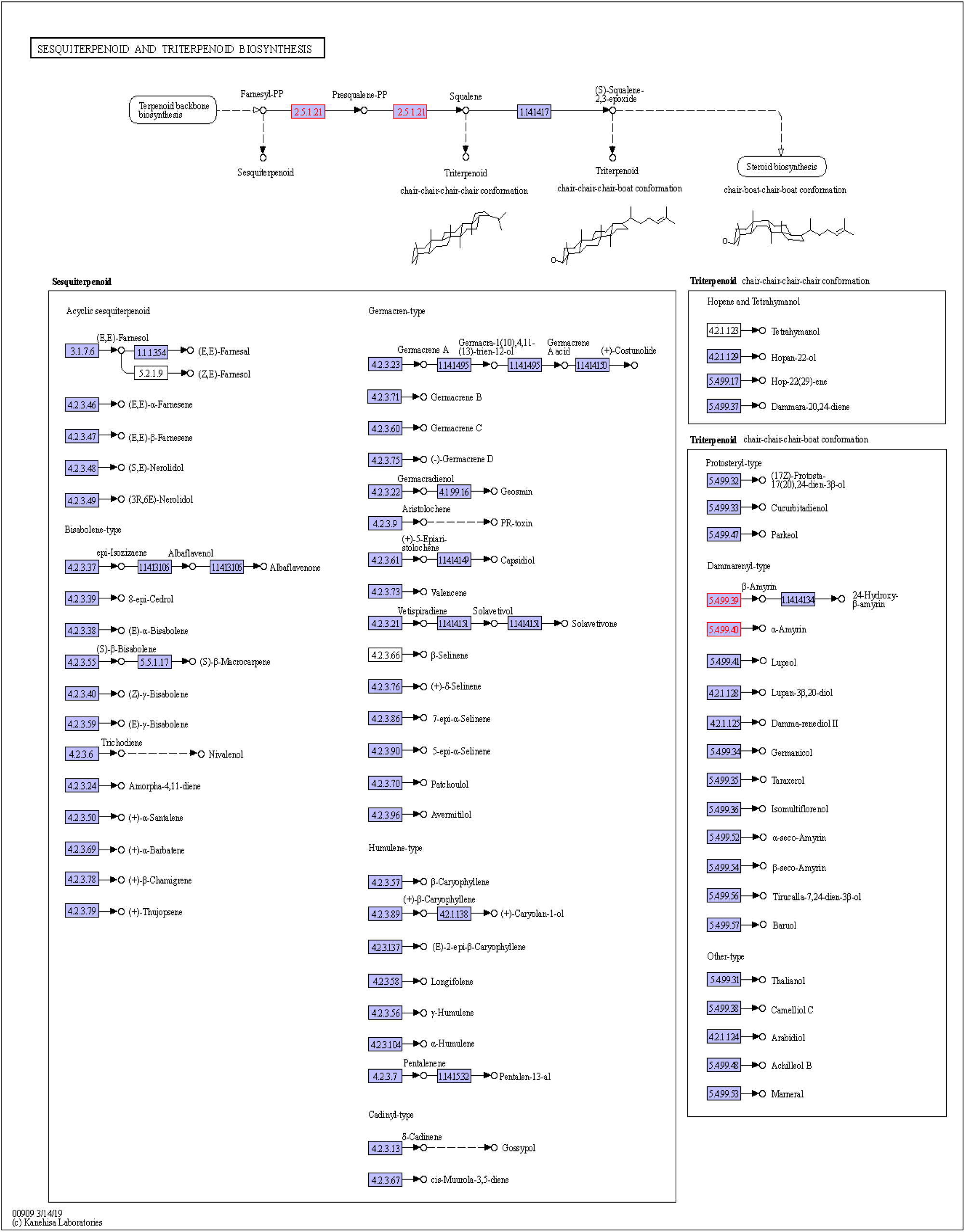
Sesquiterpenoid and triterpenoid biosynthesis pathway (ko00909) highlighted boxes are the gene found in *Taverniera cuneifolia* sequences.

**Supplementary Fig. S5:**
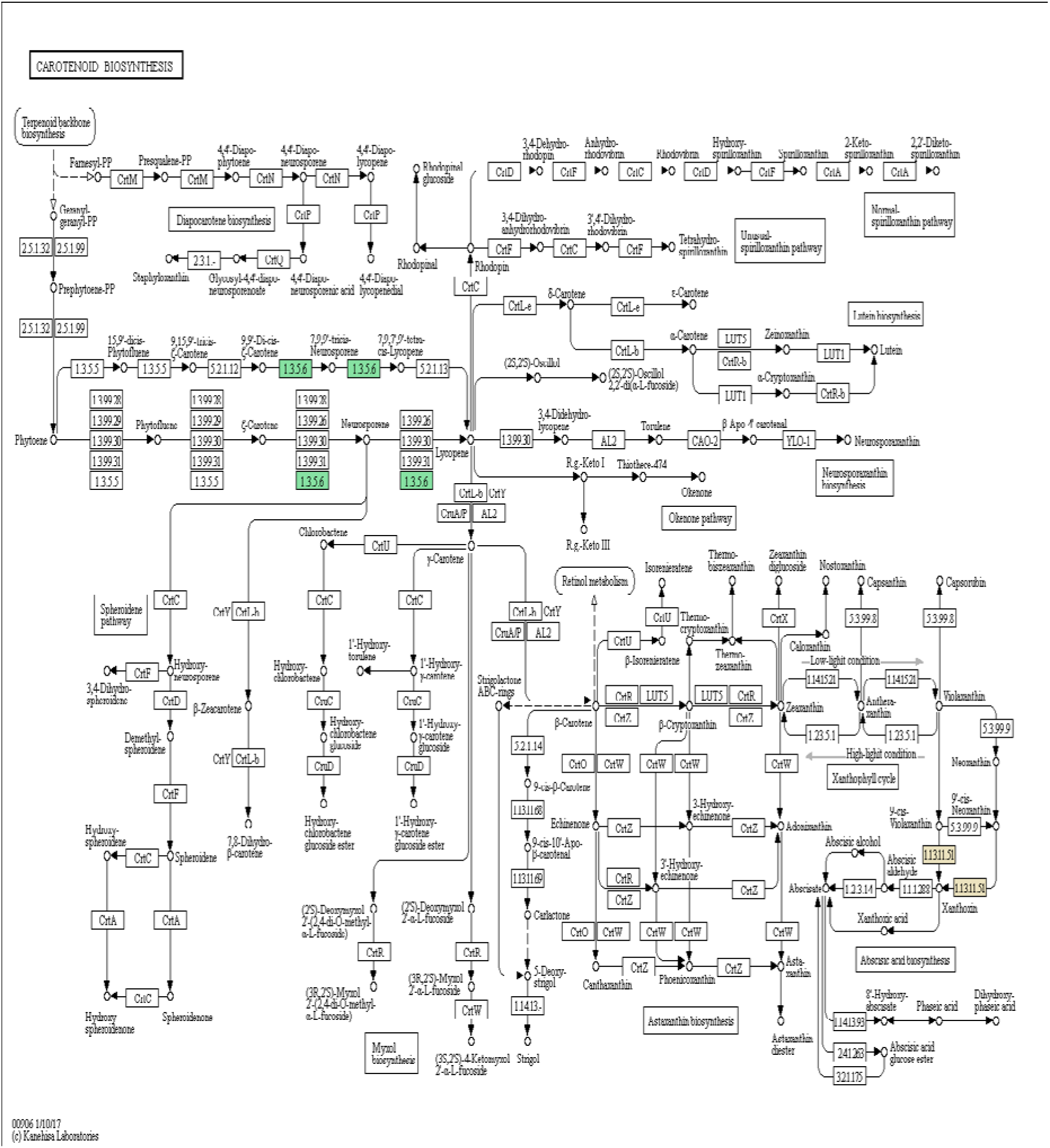
Carotenoid biosynthesis pathway (ko00906), color boxes are the gene found in *Taverniera cuneifolia* sequences.

**Supplementary Fig. S6:**
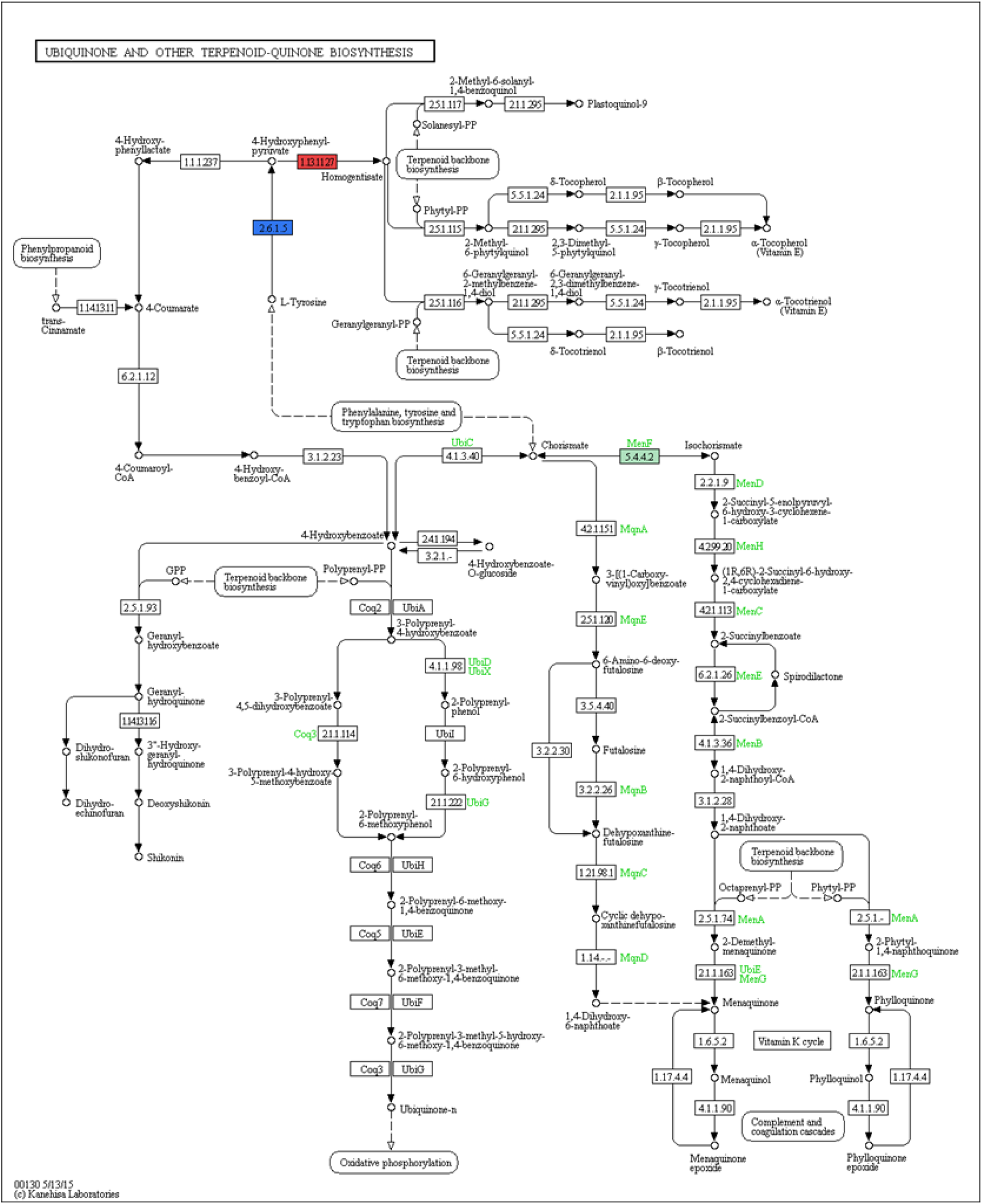
Ubiquinone and other terpenoid-quinone biosynthesis pathway (ko00130), color boxes are the gene found in *Taverniera cuneifolia* sequences.

**Supplementary Fig. S7:**
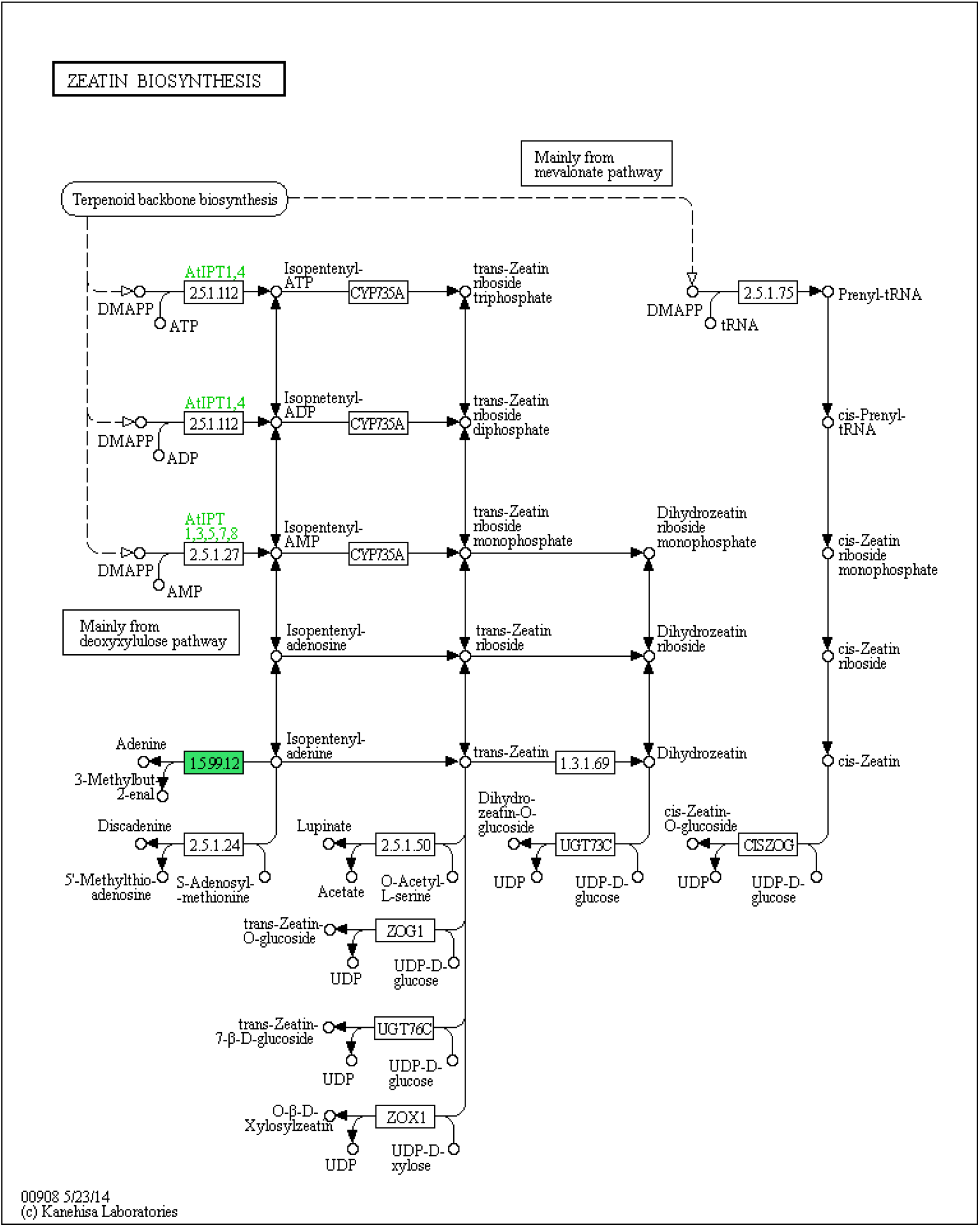
Zeatin biosynthesis pathway (ko00908), color box are the gene found in *Taverniera cuneifolia* sequences.

**Supplementary Fig. S8:**
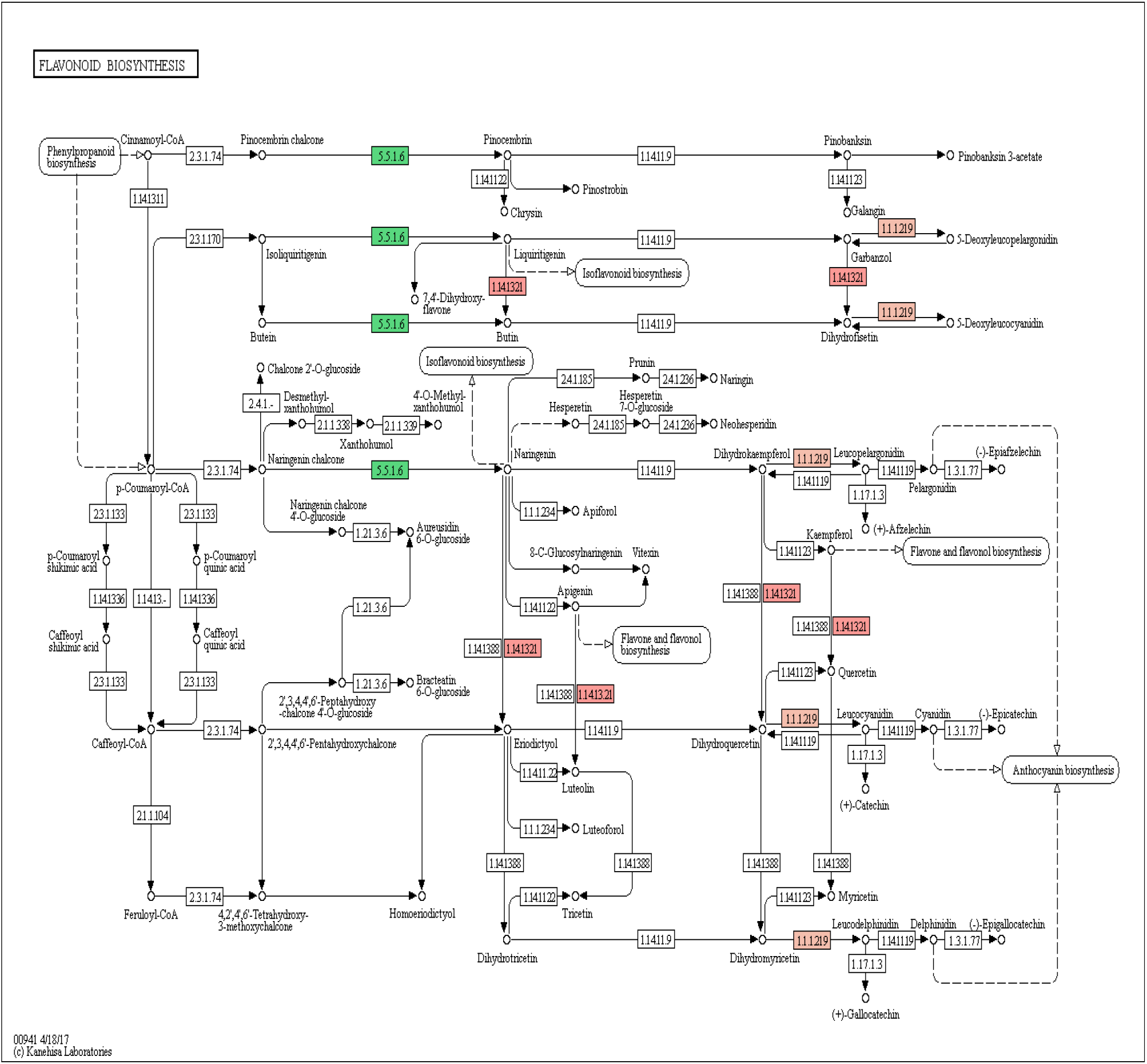
Flavonoid biosynthesis pathway (ko00941), color boxes are the gene found in *Taverniera cuneifolia* sequences.

**Supplementary Fig. S9:**
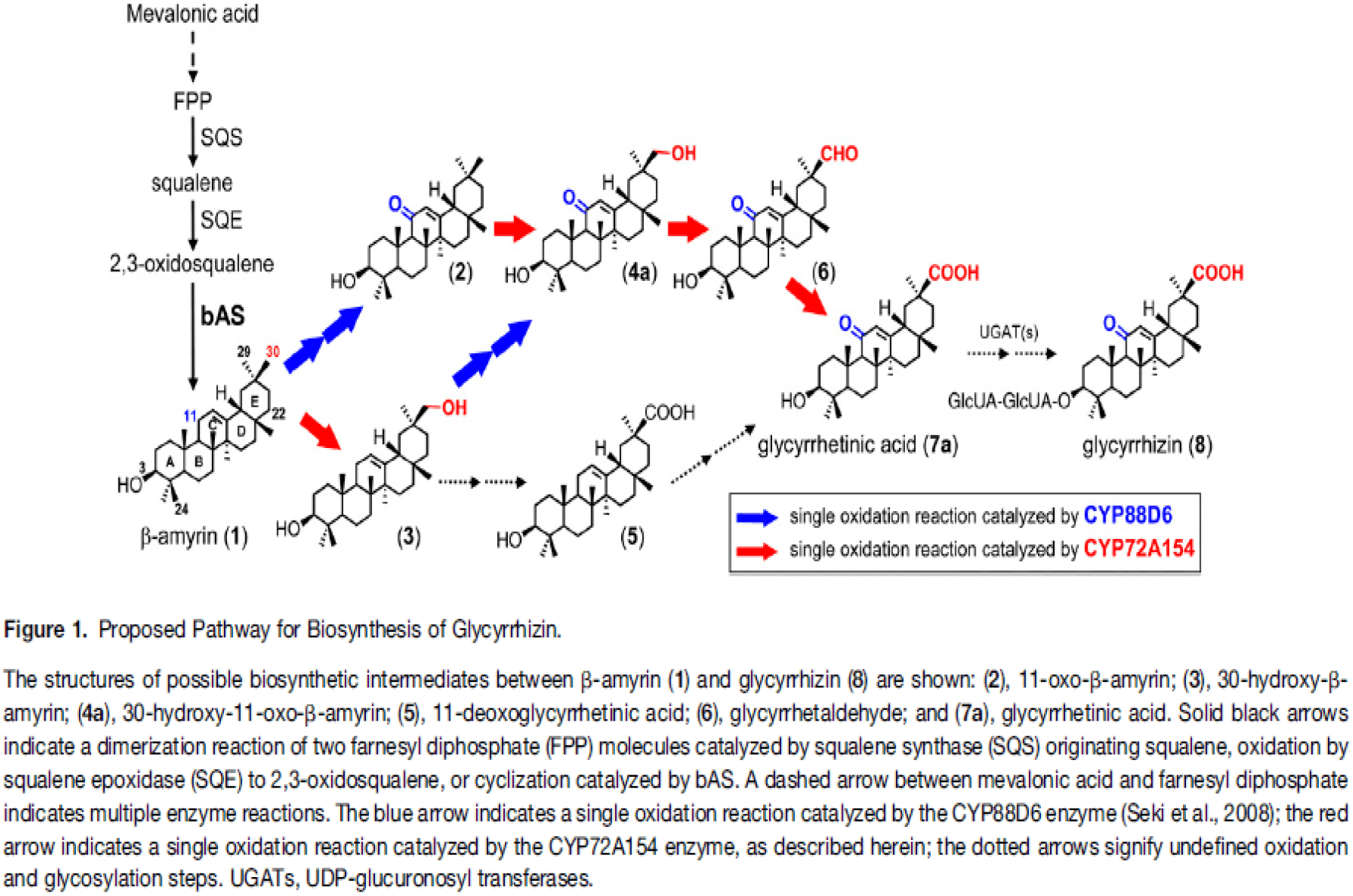
Proposed Glycyrrhizin biosynthesis pathway in Liquorice roots by seki et al 2011

